# A neural signature of reward

**DOI:** 10.1101/2022.08.23.504939

**Authors:** Luke J. Chang, Xiao Li, Kenny Nguyen, Maxwell Ranger, Yelyzaveta Begunova, Pin-Hao A. Chen, Jaime J. Castrellon, Gregory R. Samanez-Larkin, David H. Zald, Dominic S. Fareri, Mauricio R. Delgado, Livia Tomova

## Abstract

Using a neurometric approach, we identify and validate a neural signature of reward encoded in a distributed pattern of brain activity using data collected from 21 different studies (N = 2,691). Our model can discriminate between receiving rewards from punishments in completely independent data with 99% accuracy and includes weights located in regions containing a high density of D2/D3 receptors. The model exhibits strong generalizability across a range of tasks probing reward, and a high degree of specificity for reward compared to non-reward constructs. We demonstrate several applications of how this model can infer psychological states of positive affect in the absence of self report. The model is sensitive to changes in brain activity following causal manipulations of homeostatic states, can uncover individual preferences for loss-aversion, and can be used to identify positive affective experiences when watching a television show. Our results suggest that there is a shared neural signature of reward elicited across these different task contexts.

## Introduction

The survival of an organism depends upon approaching useful resources and avoiding harm. Positive and negative affective experiences associated with different potential actions can provide useful internal signals to help the agent form decision policies to successfully navigate the environment. These subjective psychological states depend upon appraising a situation (*1*, *2*) with respect to an individual’s goals, homeostatic states (*3*), and past experiences (*4*). Theories of emotion emphasize the importance of positive affect in facilitating goal attainment and reward consumption (*5*, *6*), with such states often arising following appraisals evaluating that one’s position has improved such as achieving a goal or receiving a better than expected outcome (*7*–*11*), anticipating or forecasting a positive outcome (*12*, *13*), or resolving uncertainty (*14*, *15*). The neural circuitry involved in the experience and anticipation of reward has been well studied and associated with dopaminergic neurons located in the midbrain including the ventral tegmental area and dorsal tier of the substantia nigra (*9*, *16*, *17*) that project to the nucleus accumbens in the ventral striatum (*13*, *18*) and can be gated by the medial prefrontal cortex (*19*). Though reliable activations in a network of regions including the midbrain, ventral striatum, and ventromedial prefrontal cortex (vmPFC) have been identified in meta-analyses (*20*), there is currently no pattern of brain activity that can be used to reverse infer the psychological experience of a subjective state of positive affect (*21*, *22*). Such a model would improve our understanding about how the brain processes reward and also facilitate inferences about the internal subjective experience of an individual in the absence of self-report, which could reveal insights into how humans make decisions and what happens when the positive affect system goes awry in populations struggling with mental health conditions such as depression, addiction, and mood dysregulation.

Neurometrics is concerned with assessing the reliability and validity of neural indicators of psychological states based on the fundamental principles established by the field of psychometrics (*23*–*26*). For example, there is growing evidence that patterns of brain activity measured using in vivo neuroimaging measurements (e.g., fMRI, EEG, MEG, PET, etc) can reflect information about an individual’s psychological state (*22*, *24*, *27*–*30*). These patterns can be trained using machine-learning techniques and validated using the basic principles of construct validation (*31*), in which the pattern generalizability can be evaluated to assess convergence with related psychological states elicited by different tasks and divergence from unrelated psychological states (*24*, *26*, *32*). Leveraging recent large-scale efforts to standardize neuroimaging data structures (*33*) to facilitate publicly sharing data (*34*, *35*), it is now possible to establish a so-called “nomological network”, or how a pattern of one psychological state relates to brain patterns of other psychological states (*23*, *24*, *31*). Once the reliability and validity of a brain pattern signature has been established, it can be used as an objective marker indicating the presence or intensity of a psychological state even in the absence of self-report (*36*). There is already promising evidence demonstrating the utility of this approach for negative psychological experiences such as pain (*30*), vicarious pain (*37*), negative affect (*38*), and stress (*39*). However, there has been surprisingly little work extending this approach to the domain of positive affect.

In this study, we train a multivariate brain pattern of brain activity using data collected as part of the Human Connectome Project (HCP) (*40*) in which we classify winning or losing money in a gambling task (*10*). We then demonstrate the generalizability of this pattern to a separate hold-out sample of participants also from the HCP dataset that were never included in the training of the model, and assess the convergent and divergent validity of the pattern on 17 different datasets probing a wide variety of psychological processes. Finally, we demonstrate novel applications across 3 additional datasets of how the pattern can detect changes to reward following the manipulation of homeostatic states using a deprivation study (*41*), predict individual choices in a decision-making under uncertainty context (*42*), and reveal positive affect when viewing a movie in a naturalistic context (*4*).

## Results

### Training reward brain model

We used data from the Delgado gambling task collected by the Human Connectome Project to train a brain model of reward outcomes (*40*, *43*). In this task, participants (N=490) play a card guessing game and are asked to guess if a card randomly drawn from a set of [1,9] is more or less than 5. On reward trials, participants win $1 for being correct, and on loss trials, they lose $0.50 for being incorrect (*10*). To ensure incentive compatibility, participants receive a small payment based on their performance in the game. We randomly selected 80% of the participants to serve as training data (N=392) and 20% of participants to serve as a separate hold out test dataset (N=98). A standard univariate GLM was used to perform an initial temporal data reduction, which created a map of each participant’s average brain response to reward and punishment trials. We further subtracted the subject mean out of each map and standardized the data within each image across voxels. We trained a linear Support Vector Machine (SVM) to classify reward trials from punishment trials using 5-fold cross-validation to estimate the generalizability of this model to new participants. To determine the accuracy of the model, we used forced choice tests, which compare the relative spatial similarity of each brain map to the Reward Model within the same participant using Pearson correlations and then select the map with the overall highest similarity. We perform inferences over participants by randomly permuting the order of the images for each participant 10,000 times to generate an empirical null distribution of forced choice accuracy. Using this approach, our whole-brain Reward Model was able to accurately discriminate between reward and loss maps with 98% accuracy in cross-validation, p < 0.001 (Figure 1C).

**Figure 1.**
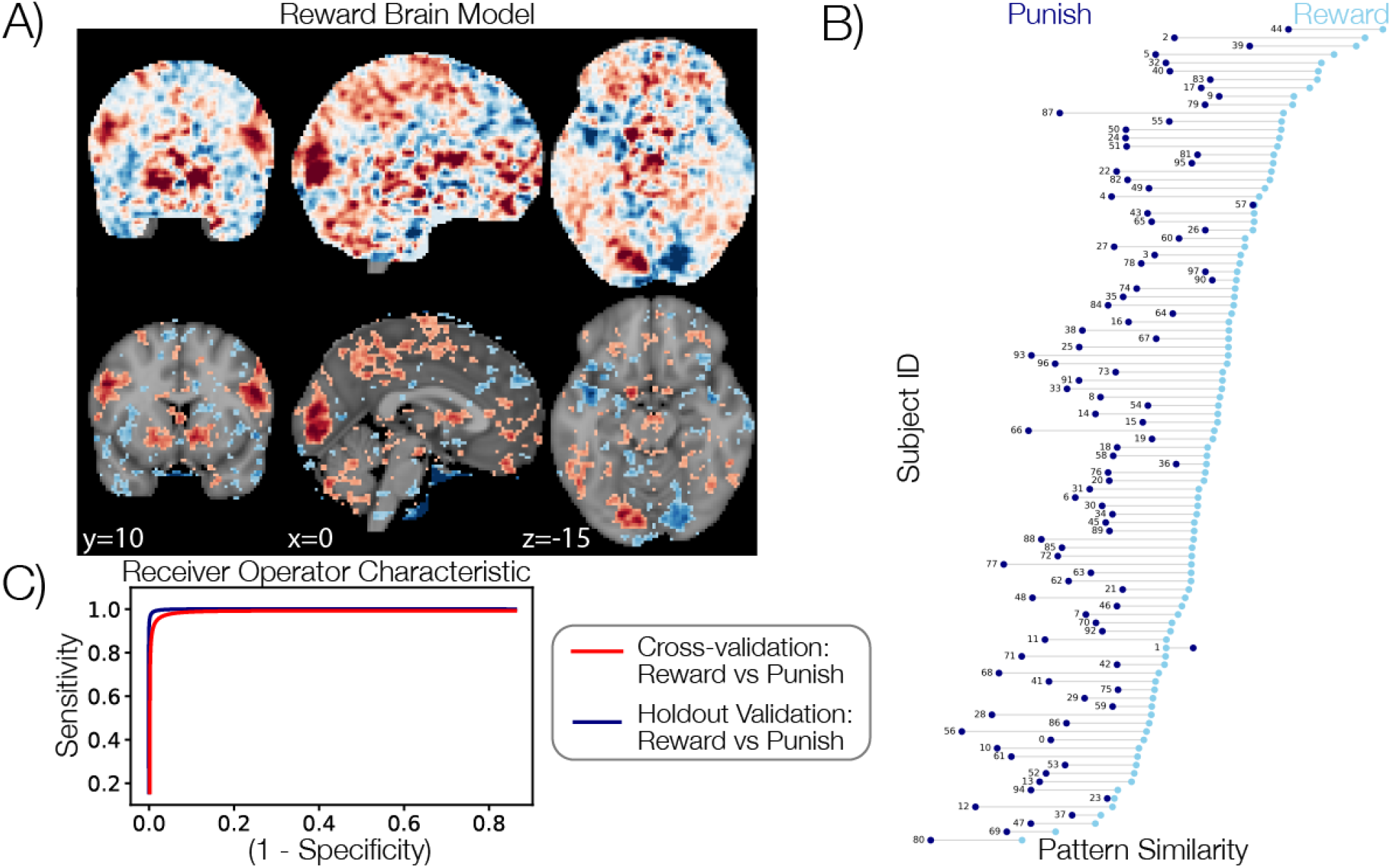
Reward Model training and validation. Panel A depicts the pattern weights for the whole-brain Reward Model using the full training dataset (N=392). Positive weights indicate an increased likelihood of classifying an image as a reward, while negative weights indicate an increased likelihood of classifying an image as a punishment. We thresholded the map at FDR q < 0.0001 to identify the most reliable weights using a parametric bootstrap procedure. Panel B depicts the results of the forced choice accuracy tests on the separate holdout dataset (N=98). The x-axis indicates the pattern similarity to the Reward Model and each line represents a single test subject. Light blue dots indicate the reward condition while dark blue dots indicate the punishment condition. Panel C depicts the receiver operator characteristic curves for testing the Reward Model in cross-validation within the training dataset (red line) and on the separate holdout dataset (navy line).

To establish the face validity of our model, we used a parametric bootstrap procedure to identify which voxels most reliably contributed to the classification, which involved retraining the model 5,000 times after randomly sampling participants with replacement and thresholding at FDR q < 0.0001 (Figure 1A). This procedure is purely for visualization and not used for spatial feature selection (*44*). The pattern of weights learned across these bootstraps exhibited a high degree of spatial consistency, r=0.93, p < 0.001 (*38*) and included expected positive weights in the ventral striatum, dopaminergic midbrain, and ventromedial prefrontal cortex (vmPFC).

Next, we trained a general whole-brain Reward Model using data from all training participants (N=392) and evaluated its generalizability to the hold-out test participants (N=98; Figure 1B). Similar to our cross-validated analyses, we found that this model was able to accurately discriminate between the reward and punishment outcomes in new participants that were not involved in training the model (forced-choice accuracy: 99%, *p* < 0.001, Figure 1B).

One potential issue using a whole brain approach is that the model may learn aspects of the psychological experience of reward that are specific to the task and do not generalize to other reward contexts. For example, outcomes in the Delgado card task are presented in the visual domain, and prior work has indicated that sensory processing may also be modulated by reward contexts (*45*–*49*). Therefore, we also trained a sparse version of the Reward Model by ablating the influence of regions outside of what is typically considered the “core reward system” using an inclusion mask created by Neurosynth (*28*) containing regions found in approximately 14,000 papers that frequently mention the word reward (see supplemental materials). Although this sparse model contained only 3% of the voxels in the whole brain model (Figure S1), it was able to accurately discriminate between reward and loss outcomes with 81% accuracy, p < 0.001 when tested in both cross-validation and also on the hold out test dataset. Thus, by virtually lesioning the model (*38*), we are able to demonstrate that the cortical regions outside of the “core reward system” improve the model performance in about 18 out of the 98 participants in the holdout test sample, which was not statistically significant using a mixed effects logistic regression Z=1.11, p = 0.27. Although both models perform well, each has its own advantages and disadvantages. The whole brain model uses information distributed throughout the entire brain and has the overall highest accuracy. In contrast, the sparse model is less accurate overall, but is also less likely to be influenced by potential confounds introduced by the task such as sensory specific effects in sensory cortices. Thus, while our primary analyses focus on the full Reward Model, there may be situations in which the sparse model can outperform the full model and so we present data from both for comprehensiveness.

### Convergent Validity

In order to demonstrate that the Reward Model is capturing the psychological state of reward, we sought to establish that it has convergent validity with reward states elicited by different tasks. We used the same forced choice accuracy validation procedure described in the model training section, which involves computing the pattern similarity between the Reward Model and the target and control conditions separately for each participant. If the model generalizes to other contexts, we predict that the target condition will be relatively more similar to the Reward Model compared to the control condition. This means that the model has conceptual overlap in the psychological processes associated with reward in the target compared to the control condition. If there is no difference between the two conditions, or if the Reward Model is more similar to the control condition, then we establish that the Reward Model does not generalize to this particular context.

First, we evaluated the spatial similarity of the Reward Model with the known spatial distribution of dopamine receptors. We used maps from the Neurosynth gene project (*50*, *51*) that are based on gene expression values derived from transcriptome-wide microarray assessments of brain tissue collected from 3,702 samples across 6 human donors provided by the Allen Human Brain Atlas (*52*). The Reward Model was most similar to the spatial distribution of gene expression values for D3 (r = 0.1), D2 (r=0.08), and D1 (r=0.04) receptors, but not D4 (r=−0.04) or D5 (−0.02) receptors (Figure 2A). Consistent with these results, we also found that the Reward Model correlated significantly with spatial patterns of D2-like receptor availability from participants (N=25) that underwent positron emission tomography scans (PET) using the high affinity D2/D3 receptor tracer [18F] fallypride, average r=0.1, p < 0.001 permuted (*53*) (Figure 2B). These results provide converging evidence that the weights learned by the Reward Model have a spatial pattern that overlaps with the spatial distribution of the density of D2/D3 receptors.

**Figure 2.**
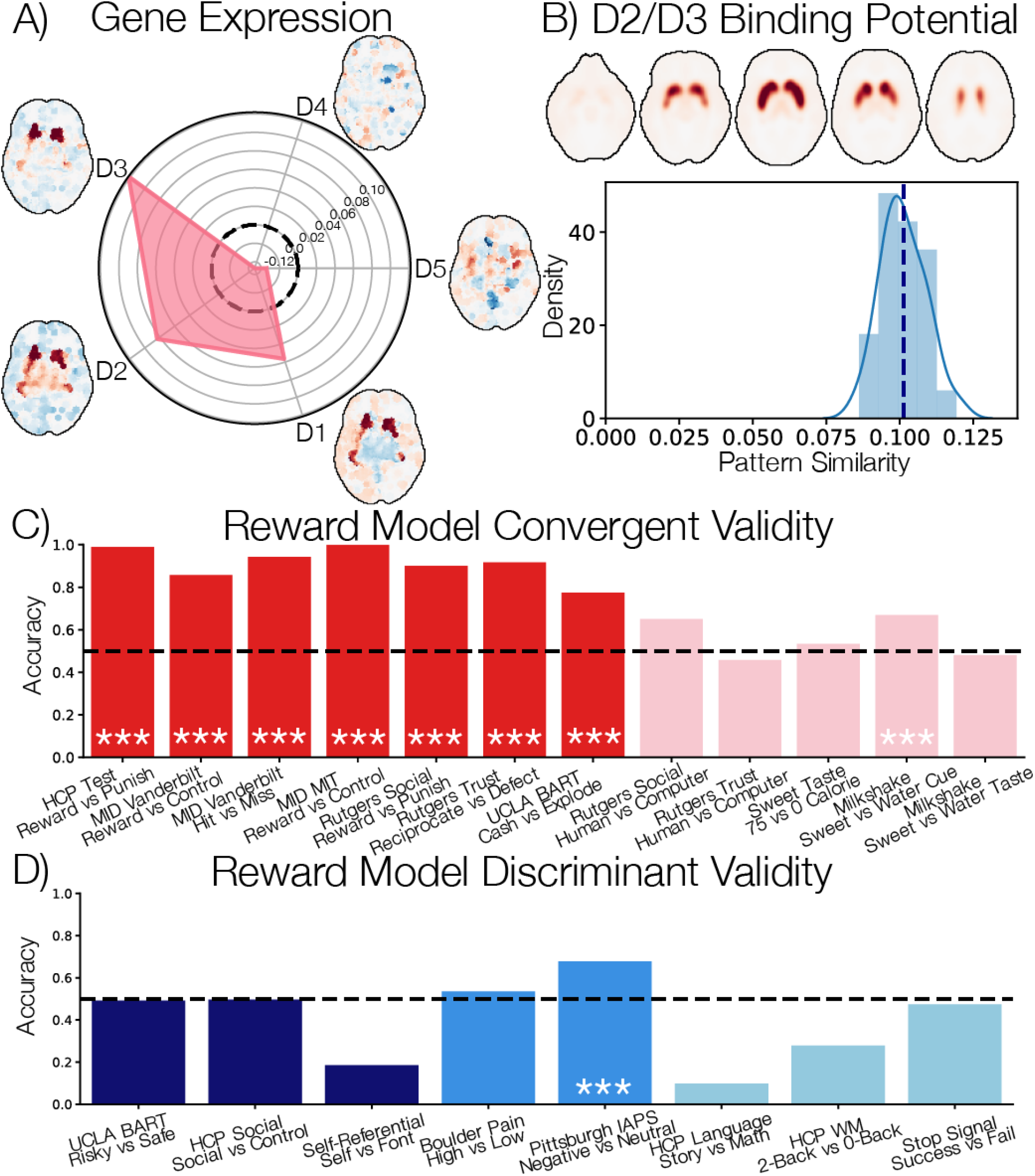
Construct Validity. A) Spatial similarity of the Reward Model with dopamine receptor gene expression derived from transcriptome-wide microarray assessments of brain tissue collected from 3,702 samples across 6 human donors provided by the Allen Human Brain Atlas. Values indicate Pearson correlation coefficients. B) Spatial similarity of the D2/D3 receptor availability from participants (N=25) that underwent positron emission tomography scans (PET) using the high affinity D2-like receptor tracer [18F]fallypride. C) Convergent validity of the Reward Model in discriminating between the reward and no reward contrasts from multiple studies. Red values are all contrasts between conditions in which participants receive a monetary outcome compared to nothing or losing money. Pink values indicate contrasts in which participants receive other types of rewards such as social or tasting sweet drinks. Statistical significance of p < 0.001 is indicated by ***. D) Divergent validity of the Reward Model in discriminating between contrasts from multiple studies. Navy values depict accuracy from risky vs safe contrasts from studies using the BART task and social and non-social contrasts. Medium blue values indicate accuracy values from negative affect studies. Light blue values indicate contrasts probing more traditional cognitive faculties such as language, working memory, and cognitive control.

Second, we assessed how well our whole-brain model of reward generalizes to other tasks designed to elicit reward. The monetary incentive delay task (MID) is among the most widely used tasks to study the anticipation and receipt of rewards (*13*). In this task, participants receive a cue indicating how much money they can win or lose if they successfully press the button before the target offset. The time viewing the cue before the target is presented provides a window into reward anticipation and the outcome period reveals how much money the participant won or lost. This task has been reliably associated with positive anticipation (*54*) and has been used as a marker of individual differences in reward processing (*55*). We found that our Reward Model successfully generalized to reward anticipation and outcomes across two variants of the MID task. Our Reward Model classified reward from neutral anticipation in a MID dataset collected at Vanderbilt (*53*) with 84% accuracy, p < 0.001, and successful from unsuccessful outcomes with 94% accuracy, p < 0.001. In addition, the model discriminated between reward anticipation from neutral anticipation with 100% accuracy, p < 0.001 in a different MID dataset collected at MIT (*41*) (Figure 2C).

Third, we examined if the Reward Model generalizes to receiving rewards in social contexts. In one study, participants play a card guessing game with shared monetary outcomes with three different partners: a friend, stranger, and computer (*56*). Our Reward Model was able to successfully discriminate between outcomes of receiving rewards compared to punishments with 90% accuracy, p < 0.001, but was unable to significantly discriminate between rewards shared with friends or strangers compared to computers, accuracy=0.65, p = 0.14. We observed a similar pattern of results in a trust game (*57*). In this study, participants made decisions to invest a $1 endowment in a relationship partner (i.e., friend or stranger), which is multiplied by a factor of 3 by the experimenter. The relationship partner then decides to either keep the $3 multiplied investment amount or return $1.5 back to the trustee (*57*). We found that our Reward Model was able to accurately discriminate between when participants learned that their relationship partner reciprocated their trust compared to defected with 81% accuracy, p = 0.001, but was unable to discriminate between when friends and strangers reciprocated compared to computers, accuracy=46%, p = 0.74 (Figure 2C). This suggests that our model successfully generalizes to social contexts involving winning and losing money, but does not appear to be sensitive to detecting with whom rewards are shared (i.e., friend compared to a computer).

Fourth, we assessed if the Reward Model generalizes to reward experiences generated from sensory modalities beyond vision. We used two independent datasets to assess how well the Reward Model could discriminate between tasting sweet drinks. In the milkshake dataset (*58*), adolescents viewed cues of glasses of a milkshake or water that signaled the impending delivery of either a 0.5ml of a chocolate milkshake or a tasteless solution. We found that the Reward Model was able to successfully discriminate between the milkshake and control visual cues, accuracy=67%, p < 0.001, but was unable to discriminate between the delivery of the milkshake compared to the tasteless solution, accuracy=48%, p = 0.71 (Figure 2C). In the sweet taste dataset, participants tasted 1ml of different drinks that were matched on sweetness using sucralose, but varied in the caloric loads by adding maltodextrin (*59*). This allowed us to assess the sensitivity of the model to experimentally manipulated carbohydrate reward. We assessed how well the Reward Model could discriminate between tasting juices matched on sweetness with 75 calories compared to 0 calories, which was shown to be associated with ventral striatal activity in the original paper (*59*). Consistent with the milkshake tests, our reward whole-brain model did not appear to generalize to tasting sweet drinks varying in caloric content, accuracy = 53%, p = 0.5 (Figure 2C). One possible explanation of these null findings is that weights in sensory cortex (e.g., occipital and insular cortex) may have been obfuscating the predictions of the model. Consistent with this hypothesis, we found that our sparse Reward Model was able to successfully discriminate between observing cues of milkshakes and water, accuracy=61%, p = 0.006, and also tasting the milkshake compared to a tasteless solution, accuracy=65%, p < 0.001 (Figure S2C). Similar results were found for the sweet taste dataset, but only approached significance, accuracy=73%, p = 0.06 (Figure 2C). These results indicate that our sparse model, which relies on areas in subcortex and the midbrain thought to be involved in core reward processing appears to be sensitive to rewards elicited from multiple sensory modalities, while our whole brain model only appears to be sensitive to detecting rewards delivered in a visual task context.

Together, this pattern of results indicates that the Reward Model exhibits strong convergent validity across a variety of tasks designed to elicit reward. The weights in the model include positive weights in regions known to contain high densities of D2/D3 dopamine receptors. The psychological processes captured by the model appear to be common in anticipating and receiving positive outcomes in a multitude of task contexts including social contexts. The whole brain model does not appear to generalize to rewards delivered in modalities beyond vision, but the sparse model can detect gustative rewards such as tasting sweet and moderately high calorie drinks (see Table S1 for full results of sparse model).

### Discriminant Validity

The next step in establishing construct validity of a model is to demonstrate discriminant validity (*31*). We hypothesized that the Reward Model should be unable to discriminate between conditions when psychological states become more distant from the experience of reward. We therefore used the same validation procedure on several other datasets that are publicly shared on OpenNeuro (*34*) and Neurovault (*35*).

First, we explored if the Reward Model was sensitive to risk and uncertainty. Making decisions requires simultaneously considering many possible uncertain consequences associated with each choice, such as the anticipated benefits and costs. Decision-making models, such as expected value, attempt to approximate the total anticipated value of each choice option by scaling each benefit/cost by the likelihood of realizing these outcomes and adding them all up. This makes it easy to figure out which choice has the overall highest expected value. Prior work has found evidence that the same regions (e.g., ventral striatum) appear to encode information pertaining both to the reward and the uncertainty or risk (*60*, *61*), while other work has found that these processes may be processed in distinct regions of the brain (e.g., ventral striatum vs insula) (*62*, *63*). Here we sought to evaluate if the Reward Model also captured aspects of the experience of uncertainty or risk using the Balloon Analog Risk task (BART), a widely used paradigm to measure risk-taking behavior (*64*, *65*). In the version of the task analyzed here, participants are presented a series of colorful (the risk condition) or achromatic balloons (the safety or control condition) and are instructed to inflate the balloons. In the risk condition, participants can choose to inflate a balloon and only receive a reward if the balloon does not explode. However, each inflation is associated with an increasing probability of explosion, and when the balloon explodes, participants do not receive a reward for that round (*64*, *65*). In contrast, in the safety condition, participants are also instructed to inflate a series of balloons, but there is no risk of the balloons exploding, nor an opportunity to receive a reward. Consistent with our results described above, we found that the Reward Model was able to accurately discriminate between reward outcomes, such as whether the participant received money from successfully inflating the balloon or lost if the balloon exploded, accuracy=77%, p < 0.001. However, the model was unable to discriminate between risky from safe decision contexts, accuracy = 0.49%, p < 0.6, indicating that the model does not appear to be sensitive to risk or uncertainty.

Second, we examined if the Reward Model could discriminate between tasks that involve social cognition. Overall, we found that the Reward Model exhibited discriminant validity by not generalizing to a variety of social-cognitive processes. The Reward Model was unable to discriminate between social vs control conditions from the HCP social cognition task in which participants viewed animate shapes that exhibited biological motion by moving in a way that conveyed social interactions compared to conditions where the same shapes moved randomly (*66*, *67*), accuracy= 50%, p = 0.59. Furthermore, the Reward Model appeared to be unrelated to self-referential processing, as it was unable to discriminate between thinking about one’s own preferences compared to a control condition in which participants were asked to make a perceptual judgment about the properties of a font (*68*, *69*), accuracy = 0.15, p = 1.

Third, we examined if the Reward Model was sensitive to arousal by testing its ability to discriminate between tasks that elicit negative affect such as pain. The Reward Model was unable to discriminate between conditions in which participants experienced high and low intensities of thermal stimulation applied to their arms or legs, accuracy=54%, p = 0.43 (*37*). However, the model was able to modestly discriminate between viewing arousing negative images compared to neutral images selected from the International Affective Pictures System (IAPS), accuracy = 67%, p < 0.001 (*38*). Interestingly, the sparse model exhibited more specificity in this context, accuracy = 43%, p = 0.93, suggesting that the whole brain model’s ability to discriminate negative arousing from neutral images may be driven by weights in visual and prefrontal cortices consistent with increases in visual attention in affectively arousing contexts (*48*, *49*, *70*).

Fourth, we assessed if our Reward Model could discriminate between tasks designed to elicit different aspects of cognition including, language, working memory, and cognitive control. We used several tasks from the Human Connectome Project. In the HCP language task, the Reward Model was unable to discriminate between reading short stories compared to completing arithmetic problems, accuracy = 10%, p = 1.0. The Reward Model was also unable to discriminate between increasing levels of working memory in an n-back task (2 back vs 0 back), accuracy = 28%, p = 1.0. Nor was it able to discriminate between levels of cognitive control using a stop signal task, accuracy = 47%, p = 0.68 (*71*).

Together, this pattern of results demonstrate specificity of the Reward Model in revealing psychological states associated with positive affect. The model finds little evidence of overlap between the psychological state of the reward captured in the Delgado card task with a variety of other psychological states spanning a range of cognitive, social, and affective processes. One notable exception is that the whole-brain model, but not the sparse Reward Model predicted a reward response when viewing negatively arousing images (see Table S1 for full results of sparse model). This indicates that the Reward Model is capturing some information that is common to both rewarding and negative experiences elicited by visual paradigms, likely reflected in visual cortex model weights.

### Applications of the Reward Model

Having now established the convergent and discriminant validity of the Reward Model, we next were interested in evaluating how we might use the model in novel ways. One of the promises of brain imaging is to reveal insights into the mind in the absence of explicit self-report. We provide several demonstrations of how this brain model might be used to understand the psychological and neural foundations of reward. First, we demonstrate that the model is sensitive to detecting changes in the rewarding value of viewing static images following causal manipulations of homeostatic states using a deprivation paradigm (*41*). Next, we show that the Reward Model can be applied to uncover participants’ subjective preferences by predicting their decisions in a mixed gambling context (*42*). Finally, we demonstrate how the model can be used to infer psychological states based purely on brain activity while participants engage in a natural viewing paradigm (*4*).

### Homeostatic State Manipulation

One interesting aspect of reward is that it is based on an individual’s subjective appraisals with respect to their goals, past experiences, and current homeostatic states. To this end, we applied the Reward Model to assess its sensitivity to changes in the subjective value of an image following a change in participants’ homeostatic states induced by a deprivation manipulation (*41*). In this study, participants were scanned across three separate sessions in which they viewed pictures of people (social condition), pictures of food (food condition), and pictures of flowers (control condition). In one scanning session, participants underwent social isolation and were alone in a room unable to communicate with another individual for 10 hours (but were able to eat as much as they wanted). In a separate scanning session, participants were food-deprived and were unable to eat for 10 hours prior to the scanning session (but were able to socialize as much as they wanted). We tested our Reward Model using the two deprivation scanning sessions and found that the Reward Model successfully discriminated between social and baseline images when socially isolated, accuracy = 87%, p < 0.001, and approached significance discriminating between social and food images, accuracy = 63% p = 0.1. When participants were deprived of food, the Reward Model successfully discriminated between food and baseline images, accuracy = 83%, p < 0.001 and also social images, accuracy=77%, p = 0.002 (Figure 3A). An even stronger test of the model is to discriminate viewing the exact same images following deprivation across sessions. Here we find that the Reward Model is unable to discriminate between social images viewed during the social isolation compared to when viewing them during the food deprivation sessions, accuracy=47%, p = 0.71. However, the model was able to successfully discriminate between viewing food images during the food deprivation session compared to the social isolation session, accuracy=70%, p = 0.022. Together, these results suggest that the Reward Model is sensitive to subtle changes in subjective valuation of images depicting social interactions or food induced by an acute deprivation.

**Figure 3.**
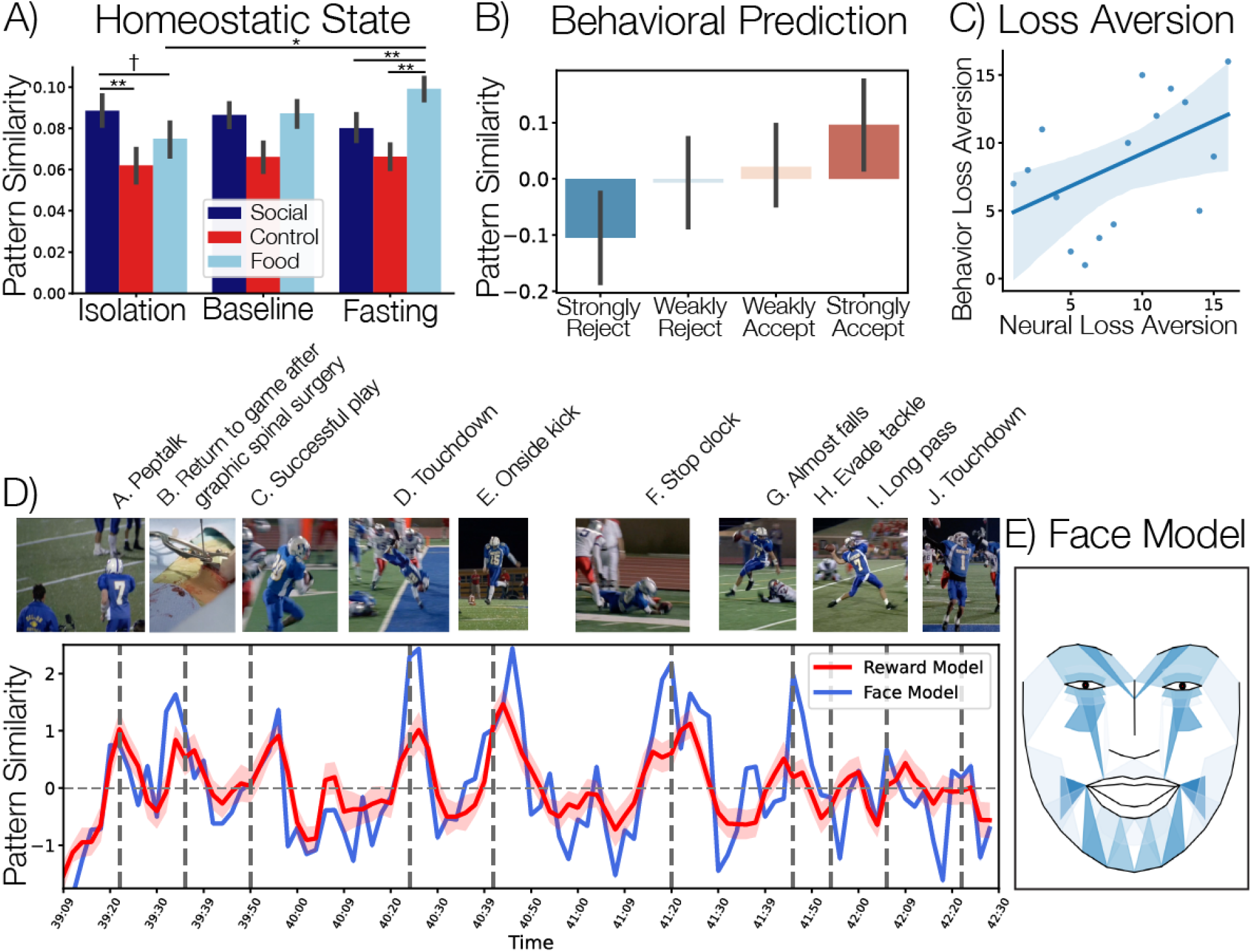
Reward Model Applications. A) the average pattern similarity of the Reward Model with each participants’ average response to viewing images across different homeostatic state manipulations (41). B) the average pattern similarity of the Reward Model to viewing mixed gambles conditioning on participants’ individual decisions to accept or reject the gamble (42). C) Relationship between λ loss-aversion parameter estimated from both participants’ decisions and their neural activity predicted by Reward Model. Units are in ranks to correspond to our Spearman ρ analysis. D) Red line depicts the average Reward Model pattern similarity to passively viewing the final football scene from the pilot episode of Friday Night Lights (4) error bars reflect 95% confidence intervials. Dotted dark gray lines indicate the outcome of each play. Blue line depicts the average predictions of the cross-validated facial expression model trained to predict the Reward Model pattern similarity. Screenshots of each scene are copyright of NBCUniversal, LLC. E) Facial expression model that best predicts the pattern similarity of the Reward Model from the fMRI study. The color intensities indicate how much each facial action unit (AU) contributes to the time-series prediction. Weights are normalized between [0,1] for display purposes.

### Estimating Loss Aversion

Next, we sought to test the ability of the Reward Model to reveal an individual’s subjective preferences in the context of a mixed gamble task. We used an open dataset, in which participants (n=16) were asked to decide whether to accept or reject mixed gambles that varied in potential gains or losses (*42*). Using a mixed effects logistic regression, we successfully replicated the behavioral results reported in the original paper. Participants were more likely to accept a gamble as the potential gain increased, *β*_Gain_ = 0.43, se = 0.08, p < 0.001, and less likely to accept as the potential loss increased, *β*_Loss_ = −0.72, se = 0.07, p < 0.001. Next, we were interested if we could find similar results derived purely from brain activity using our Reward Model. For each participant, we estimated a single trial first-level GLM to generate brain maps to each gamble, and computed the spatial similarity with our Reward Model to assess the predicted reward values to each trial. We used a mixed effects regression to estimate the influence of the potential gain and loss amounts on the predicted reward responses and observed that independently varying losses (*β*_Loss_ = −0.02, se = 0.004, p < 0.001), but not necessarily gains (*β*_Gain_ = 0.003, se = 0.002, p = 0.22) significantly predicted neural reward responses (Figure 3B). Though our Reward Model was not able to reliably independently parse gain from loss signals (perhaps because it was originally trained to discriminate between rewards and losses), the model was able to significantly predict participants’ trial-to-trial decisions to accept or reject a gamble, *β* = 6.87, se = 2.07, p < 0.001 (Figure 3B). Moreover, a forced choice accuracy test averaging over trials within each participant revealed that the Reward Model was highly accurate in classifying participants’ decisions to accept or reject the gamble (accuracy=94%, p < 0.001). Finally, we assessed how well we could estimate a participant’s subjective loss-aversion preferences λ using a linear model of prospect theory (*42*), where *β*_Loss_ and *β*_Gain_ are estimated from a mixed effects regression, *P*(*Accept*) = *β*_*Intercept*_ + *β*_*Gain*_ + *β*_*Loss*_, and 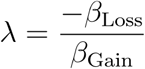. We observed a significant positive relationship between participants’ subjective loss-aversion preferences estimated from their decisions and also purely from their brain activity based on the trial-to-trial predictions of the Reward Model, *ρ* = 0.48, p = 0.03 permuted (Figure 3C).

### Uncertainty resolution during movie watching

As a final application of the Reward Model, we sought to assess how well it could uncover an individual’s subjective experience during a passive viewing context. While all of the previous tests used tasks designed to probe a specific psychological process, movie viewing tasks vary in many dimensions simultaneously and provide a rich testbed for evaluating the generalizability of a model. Unfortunately, it is difficult to annotate a movie on all psychological dimensions, and participants also tend to display more individual variability compared to traditional tasks (*4*, *72*, *73*). In this dataset, participants (N=35) watched the 45-minute pilot episode of a character driven television drama, *Friday Night Lights* (*4*). We applied the Reward Model to participants’ timeseries while they watched the movie and specifically examined the predictions of the model to a sequence of dramatized plays in a 3.5 minute clip at the climax of the episode, in which the backup quarterback leads the team to an unexpected victory. This sequence of scenes provides a particularly interesting test of the model as there are multiple events that elicit a high level of uncertainty (e.g., will the receiver catch the ball?) before the outcome is revealed (Figure 3D). In this short clip, at the end of the episode, the backup quarterback receives a pep talk from his coach during a time out (A), then makes a successful handoff to a running back (C), which then leads to scoring a touchdown (D). With only minutes left in the game, the team makes a risky onside kick, and is able to successfully recover the ball in a turnover (E). With less than a minute left, the backup quarterback makes a successful pass and the receiver steps out of bounds to stop the clock, saving precious seconds (F). In the final play of the game, the quarterback almost falls following the snap (G), then successfully evades a tackle (H), and throws a very long pass (I), which is eventually caught and leads to the game winning touchdown (J). A common technique to assess the reliability of a signal across participants in naturalistic designs is to compute the intersubject correlation (*74*, *75*). Here, we examine the temporal dynamics of the Reward Model pattern similarity across participants and observed a high degree of consistency during this short sequence (temporal ISC = 0.31, 95% CI = 0.29-0.36, p < 0.001), which was comparable to the temporal ISC across the entire episode (ISC = 0.35, 95% CI=0.32-0.40, p < 0.001) and also to the amount of temporal synchronization we observed in early visual cortex across participants (ISC = 0.33, 95% CI=0.28-0.40, p < 0.001), which is a region that tends to exhibit the highest degree of intersubject synchronization across the entire brain (*4*, *74*). These results indicate that the Reward Model is capturing a reliable signal across participants corresponding to the resolution of the uncertainty of each event consistent with the protagonists triumphing in the game. However, this does not necessarily mean that the model is capturing psychological processes associated with reward, which would require additional information about how the participants were feeling while the narrative unfolded.

To gain further insight into the psychological processes being captured by the model, we used an additional behavioral dataset to decode the affective experience captured by the Reward Model. In this dataset, a different sample of participants (N=20) watched the same television episode while their facial expressions were video recorded (*4*, *76*). We used a computer vision algorithm to automatically identify the temporal dynamics of 20 facial action units (AUs), which represent a standardized system to describe the intensity of facial movements (*77*, *78*) at each frame of the video. We then used the normalized averaged time series of each AU across this sample to predict the temporal dynamics of the Reward Model pattern similarity to the fMRI data described above during the sequence of scenes using linear regression with 5-fold cross-validation. Overall, we found that the face model was able to accurately predict the Reward Model pattern response during this sequence of football plays in new participants, mean *ρ* = 0.39, sd = 0.12, p < 0.001 permuted. Consistent with our hypotheses, the learned face expression reflects a behavioral demonstration of a positive affective experience of anticipation (Figure 3E) with the largest increases in AU14 (Buccinator, dimpler), AU9 (Levator labii superioris alaquae nasi, nose wrinkler), and AU25 (Depressor Labii, lips parting), demonstrating the potential of these models for supplanting self-report.

**Table 1.**
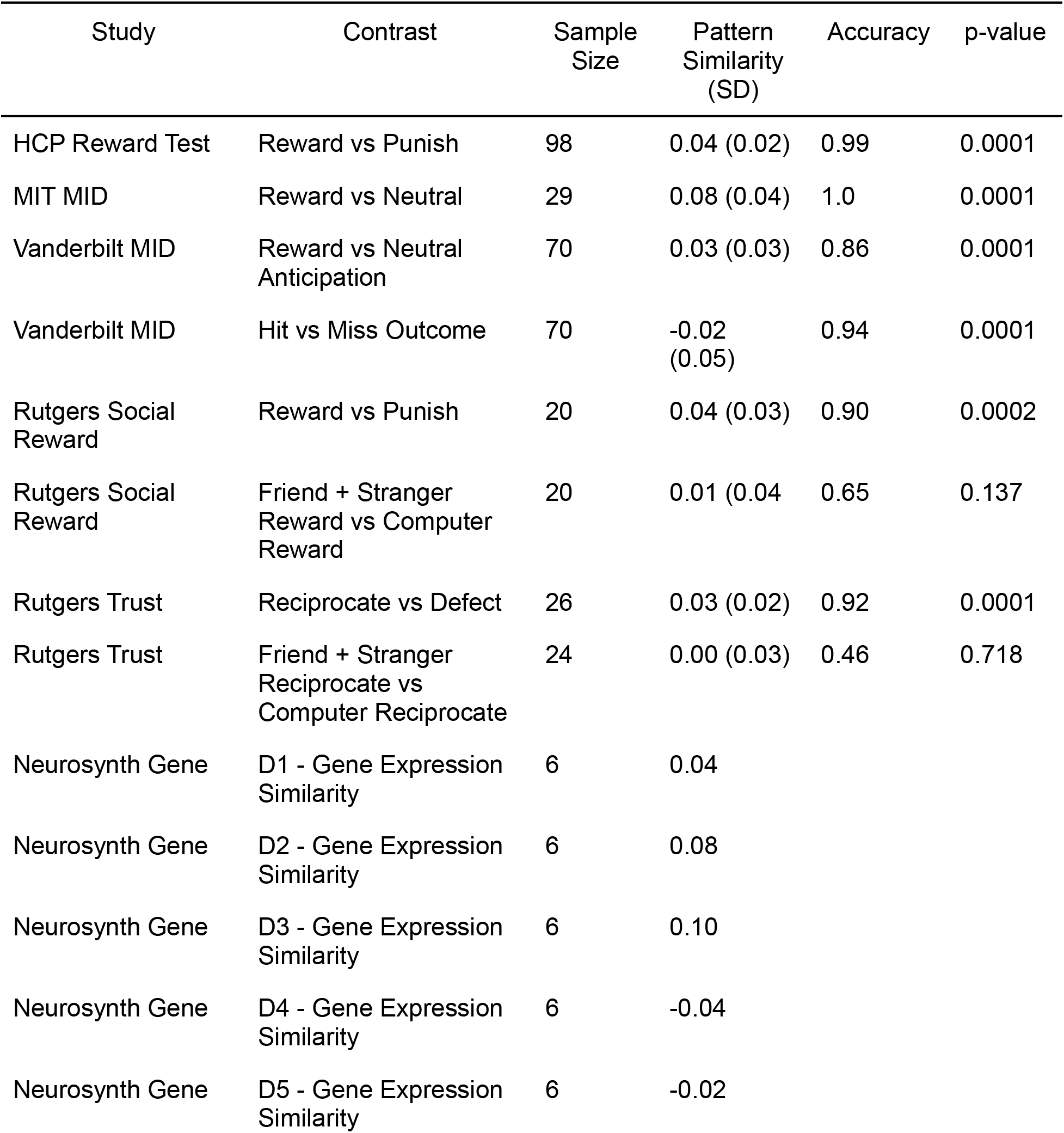
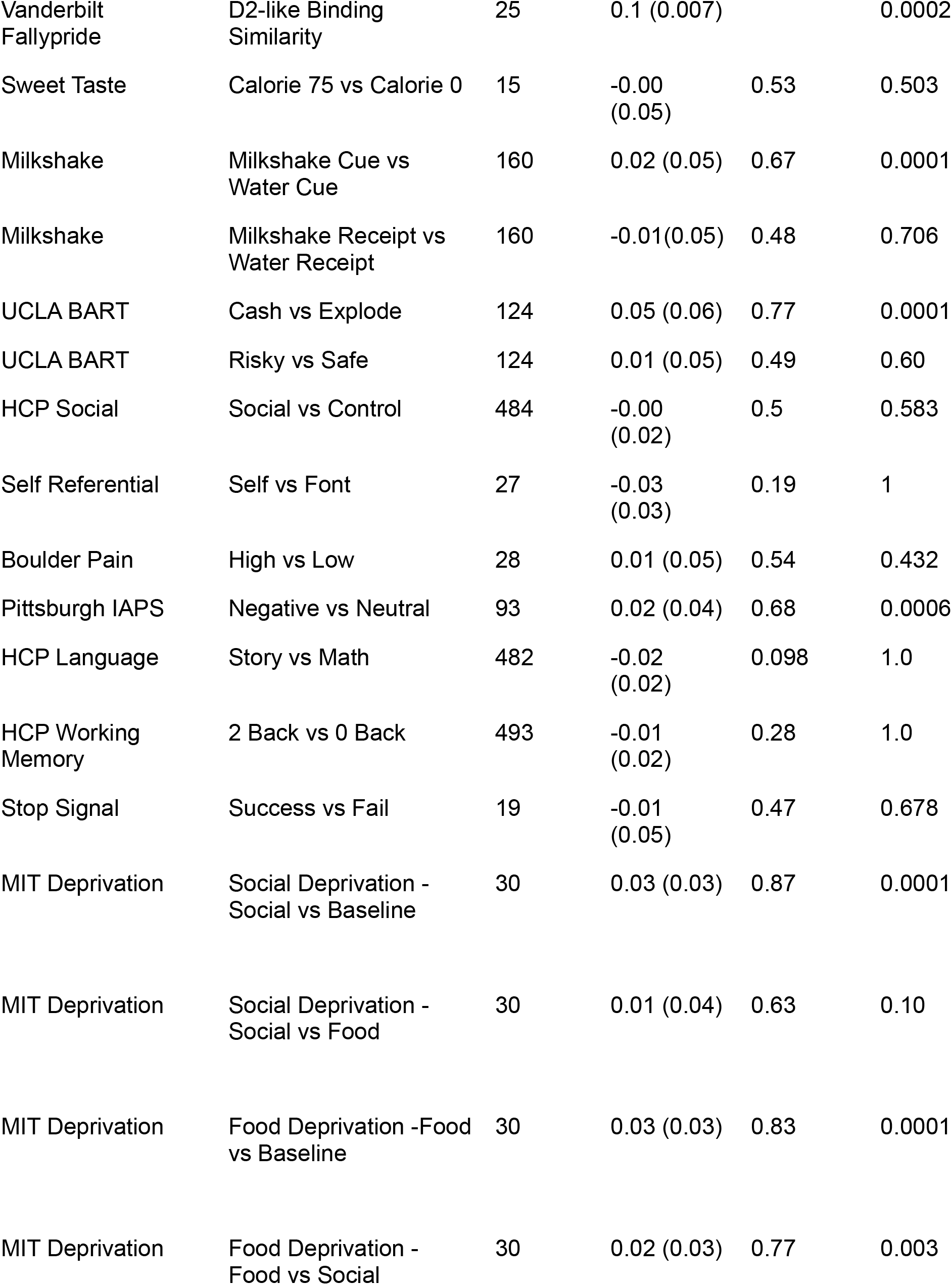
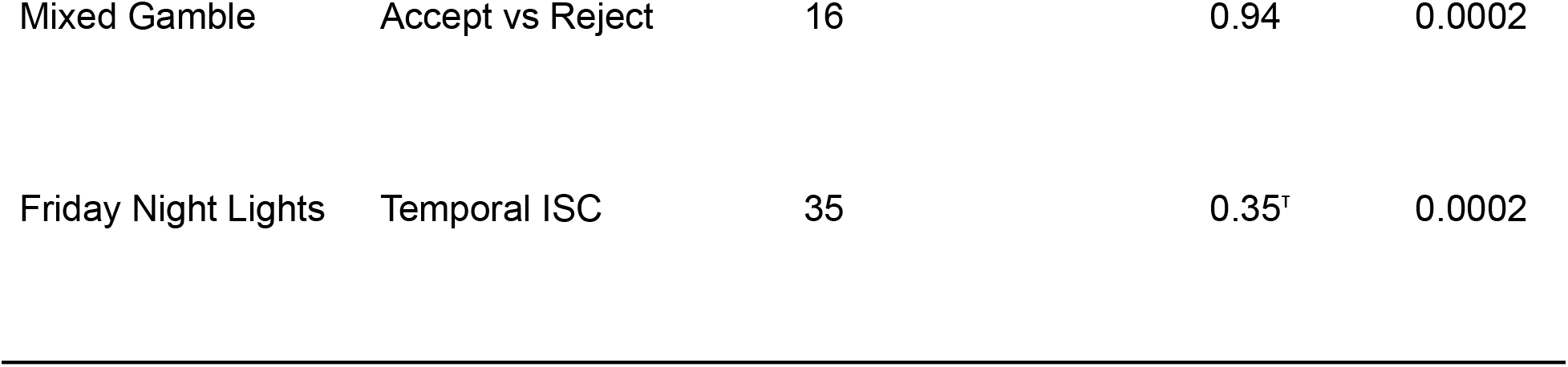
Convergent and Discriminant Validity of Reward Signature. ^T^indicates temporal intersubject correlation (ISC). See Table S1 for Sparse Reward model results

## Discussion

In this study, we develop and validate a neural signature of reward using whole-brain multivariate pattern analysis using publicly shared data from 21 different studies (combined N = 2,691 participants). We find that a linear model trained on a large gambling dataset collected as part of the Human Connectome Project (*40*) is able to discriminate between outcomes where new participants won money compared to lost money with 99% accuracy (*10*). This multivariate pattern exhibits strong generalizability across participants, scanners, and tasks probing reward. Moreover, the weights that contribute most to the model's predictions are located in regions that have previously been associated with a high density of D2/D3 receptors suggesting a possible connection to dopaminergic activity. The Reward Model also shows a high degree of specificity to reward relative to psychological constructs unrelated to reward across a range of cognitive and affect processes such as risk, working memory, cognitive control, language, social cognition, and pain. We further demonstrate several potential applications of how this model might be used as a method to infer psychological states of positive affect in the absence of self report. The model is sensitive to subtle changes in the positive affective experience following a causal manipulation of homeostatic states using a deprivation paradigm. The model is also able to accurately predict participants’ decisions in a mixed-gambling task and can be used to estimate individual participants’ subjective loss-aversion preferences. Finally, the model can be used to identify positive affective experiences when watching a television show in the scanner.

Using a neurometric approach to evaluate the construct validity of this model provides unique insights into the construct of reward. Though the concept of reward is central to many disciplines ranging from economics to neuroscience, its ubiquity belies its complexity. Our model was designed to capture the psychological experience of reward elicited in the context of winning a gamble and measured using BOLD fMRI. It generalizes extremely well to other tasks that share a similar structure that includes a visual presentation of states of the task, and the resolution of uncertainty with a positive outcome (*7*–*11*). These positive outcomes can include winning money, finding out that you were correct about something, or watching your favorite team win a game. The model also seems to capture the psychological state of anticipating an imminent favorable outcome (*12*, *13*) such as winning a gamble, or knowing you are about to get a sip of a tasty drink. The reward processes captured by the model are not inherently tied to a particular state of the world or outcome, but rather to an individual’s appraisal of the subjective meaning of the event at a specific place and time. These appraisals are malleable and can be changed when the agent pursues different goals, which can be altered by fluctuations in internal homeostatic states (*4*). We find that viewing stimuli associated with basic needs such as food or social interactions is predicted by the model to be more rewarding when you have been deprived of these needs (*3*, *41*). Furthermore, individuals do not necessarily value the same things. For example, variations in individual sensitivity to prospects of gains relative to losses, a phenomenon known as loss aversion (*42*, *79*), has traditionally been identified based on decision behavior. Here we show that loss aversion preferences can be estimated purely from patterns of brain activity when contemplating a gamble using our Reward Model.

Interestingly, the full whole-brain Reward Model did not perform as well in contexts that dramatically differed from the original training context. Tasting sweet drinks may be perceived as rewarding (*58*, *59*), but our model was confused when the experience was elicited via gustative rather than visual input. It is possible that these other areas are providing information that is not directly related to reward. For example, we also found that the whole-brain model predicted that viewing negative aversive images was rewarding, but the sparse model did not make this mistake. One possible explanation is that inputs from sensory and prefrontal cortex may be capturing processes associated with visual attention that are present in both positive and negative affective states (*45*, *46*, *48*, *49*, *70*). Another possibility is that these connections with other cortical regions may contextualize the rewarding experience, which could create a wider range of positive affective experiences than can be captured by the sparse Reward Model. This suggests that ongoing arguments in the field about whether the brain represents different types of value signals as a common currency (*80*, *81*) may not be able to be resolved from simply examining overlapping activity in the striatum; interactions between regions may contribute additional nuance. Moreover, it is highly likely that BOLD fMRI itself does not have enough temporal or spatial sensitivity to detect such differences. For example, there is evidence that distinct neurons in adjacent regions of the striatum in non-human primates independently track social and non-social rewards (*82*), rewards to self and others (*83*), and approach and avoidance motivation (*84*).

Although this paper provides a wide range of tests supporting the sensitivity, specificity, and generalizability of our model, there are a number of limitations that are important to acknowledge. First, these tasks only represent a fraction of the types of rewarding experiences that are likely possible. These tasks are easy to run in the scanning environment, but cover a very limited scope of possible rewarding experiences. Moreover, the structure of our validation procedure precluded more nuanced methods of inferring rewarding experiences that might leverage individual differences or subjective experiences of reward measured via self-report. Though we used popular tasks that have been previously shown to elicit reliable findings, none have undergone rigorous neurometric testing to assess the overlap in the potential psychological processes elicited. Second, we do not view this model as the end goal, but rather the beginning of a prolonged process of refining the model and continuing to assess its sensitivity and specificity as more datasets become available. These results merely provide an initial benchmark, which we hope will be improved upon by other groups. We made numerous assumptions to make this work tractable such as assuming linearity and that information could be contained in averaged activity within independent voxels distributed throughout the brain. Of course different feature selection approaches, and looking at interactions between regions, along with assuming more complicated nonlinear functions will likely surpass our model in terms of its generalizability, but also in helping us improve our understanding of how the brain represents reward. For example, we trained our model using a single task using a linear shallow-learning approach. Combining all of the reward contexts and more powerful deep-learning models will likely facilitate learning richer representations of reward.

One of the promises of in vivo neuroimaging is to accelerate learning about the functional organization of the brain and the neurobiological basis of psychological processes. In this paper, we develop and validate a model of positive affect based purely on patterns of brain activity that can objectively quantify subjective experiences of reward in the absence of self-report. This model may be of broad interest to facilitate new directions in studying social and affective processes (*24*). For example, does the neural representation of reward change across development? Perhaps patterns of different rewarding experiences are more distinct earlier and become more similar as we grow older? In addition, this model may provide further insights into the nature of impairment in the multiple psychiatric disorders in which alterations in reward processes figure prominently. In this context, individual deviations from a “canonical” Reward Model could provide useful treatment planning information. For example, measuring an individual's reward signature over time could provide a means of tracking the effect of clinical interventions and should have an advantage in that it may be less sensitive to placebo effects and participant bias than conscious self-reports of mood. We openly share both the whole-brain and the sparse Reward Models and recommend that these models be used interchangeably depending on the researcher’s goals. This paper builds on an enormous amount of work over the past thirty years that required refining imaging acquisition, preprocessing, and analytic techniques. In addition, it leverages large concerted efforts to standardize how neuroimaging data is represented and reported (*33*) and also open source tools to share and access this work (*28*, *35*, *85*), as well as a strong commitment by the broader neuroimaging community to curate and openly share their data to facilitate secondary data analyses (*86*). We hope that this model of reward and the neurometric approach outlined in the paper will in turn help facilitate new ways to study the brain to accelerate the pace of discovery.

## Methods

### Datasets

#### HCP Delgado Gambling Task

The primary dataset we used to train the Reward Model is the Delgado gambling task (*10*) collected as part of the Human Connectome Project (*40*, *43*). In this task, participants (N=491, mean age = 29.24, 59% female) play a card guessing game and are asked to guess if a card randomly drawn from a set of [1,9] is more or less than 5. On reward trials, participants win $1 for being correct, and on loss trials, they lose $0.50 for being incorrect (*10*). To ensure incentive compatibility, participants receive a small payment for what they believe is based on their performance in the game. All participants provided informed consent in accordance with protocols approved at Washington University. The HCP acquired fMRI data on a Siemens Skyra 3T scanner using a gradient-echo echo planar imaging (EPI) sequence with a multiband acceleration factor of 8 (TR=720ms, TE=33.1ms, flip angle=52°, FOV 208 × 180mm, matrix size=104 × 90, voxel size= 2 × 2 × 2). Data were preprocessed according to the minimal preprocessing HCP fMRIVolume pipeline (*87*), which includes removing spatial distortions via gradient unwarping and fieldmap-based distortion correction, realignment, and spatially normalizing the data to MNI space using a brain-boundary-based registration and nonlinear transformation implemented via FSL. In addition, the data were temporally filtered using a Gaussian-weighted linear highpass filter with a cutoff of 200s, and prewhitened using FILM to correct for temporal autocorrelation. First level models were run using FSL and included boxcar regressors for each experimental condition convolved with a double gamma canonical hemodynamic response function (HRF), and temporal derivatives of these regressors to account for variability in HRF delays. Data were smoothed with a 4mm Gaussian kernel. The beta estimates for the average response to each condition were accessed via the OpenAccess AWS S3 bucket provided by the HCP (https://www.humanconnectome.org/). We note that in the original analyses reported by the HCP, there were no regions with activations greater than a z-threshold of 1.96 in at least 50% of the sample in the reward vs punishment comparison, indicating substantial individual heterogeneity in this dataset and low temporal signal to noise ratio (tSNR) in the striatum and orbitofrontal cortex (OFC) (*40*).

#### MIT Monetary Incentive Delay

In this study, participants (N = 29, mean age=26.9, sd=5.8, females=76%) completed a monetary incentive delay (MID) task as a functional localizer during the baseline condition of a social isolation experiment (*41*). The task was adapted from (*88*) and all participants provided informed consent in accordance with an approved protocol by the MIT Institutional Review Board. Before beginning the task, participants memorized a set of 5 images depicting abstract art (all images taken from the free stock pictures site (https://www.pexels.com/)). During the task, the abstract art images served as cues to the condition of the current trial. The task had two conditions: a reward/loss condition (*reward*) in which participants could earn or lose money depending on whether their responses were correct and fast enough, and a non-reward condition (*non-reward*) in which participants always received $0 regardless of their response. Each trial started with an abstract art image. The previously memorized (*familiar*) images indicated a *non-reward* trial. Abstract art images that were not previously observed (*novel*) indicated a *reward* trial. After the cue, participants saw a number between 1-9 (excluding 5) for 100ms on the screen. Their task was to press an assigned button indicating whether the number is below or above 5 as fast as possible. Initially, correct responses were required in less than 500ms; after 10 consecutive correct answers, this window was reduced to 400ms. After they pressed the button, participants saw the outcome indicating whether they won $1 (*reward* trial, correct response, fast enough), lost $0.20 (*reward* trial, wrong response or too slow), or received $0 (*non-reward* trial). In total, participants played 80 trials (40 trials per condition) and the duration of the task was approximately 10 minutes. In this dataset, we were only able to examine the anticipated reward phase as the motor response and outcome phases were not designed to be temporally separable. Data were acquired on a Siemens Prisma (TR = 2,000 ms, TE = 30 ms, FOV = 210 mm, 70 × 70 matrix, yielding a voxel size of 3 × 3 × 3 mm^3^) acquired as a partial-head volume in an anteroposterior phase-encoding direction using interleaved slices and were accessed from OpenNeuro (*85*) (https://openneuro.org/datasets/ds003242/versions/1.0.0). Data were preprocessed using fMRIPREP (*89*) and smoothed using a 6mm FWHM Gaussian filter. First level single-trial models were run using nltools (*90*). Each trial was convolved with a double gamma HRF and additional covariates included a linear trend, the effects of motion estimated during the realignment step using an expanded set of 24 motion parameters (six demeaned realignment parameters, their squares, their derivatives, and their squared derivatives), motion spikes between successive TRs, and global signal-intensity spikes greater than three SDs above the mean intensity between successive TRs, and a high pass filter of 120s.

#### Vanderbilt Monetary Incentive Delay

In this study, participants (N = 70, mean age=38.8, sd=15.6, females=51.4%) completed a probabilistic variant of the monetary incentive delay task (*62*) designed to assess sensitivity to varying reward magnitudes and probabilities (expected values). All participants provided informed consent and this study was approved by the IRB at Vanderbilt University. During the task, participants were first shown an explicit cue (2000 ms) that indicated one of 12 possible combinations of reward magnitudes in US dollars ($0, $1, $2, or $3) and probabilities (20%, 50%, or 80%). Following the cue, a brief fixation cross was presented for a variable amount of time (2000 to 2500 ms) followed by a target (100 to 400 ms) during which the participant was shown a white star and was required to quickly press a key on a response box. The target screen was followed by another brief fixation cross (2000 ms minus target duration) before participants were presented with feedback (2000 ms) that indicated whether their key response was quick enough. As in Samanez-Larkin et al., (*91*), the hit and miss rate for individual participants was manipulated by altering the average duration of the target with an adaptive timing algorithm that was originally set to the individual’s mean reaction time in a pre-scan practice, and then followed their performance across the scanned blocks, such that the individual would successfully hit the target on approximately 66% of the trials for each cue type. Trials were separated by a variable intertrial interval (2000 to 12000 ms). The task was divided into 3 runs with 42 trials in each run. At the end of each run, participants were shown the amount of money earned. The task was incentive compatible; participants were told that they would be paid 20% of their total earnings from the task. Actual total earnings from participants varied between $14 and $31 (M = $21.64, SD = $2.27). Data were acquired on a 3T Philips Intera Achieva scanner using a T2*-weighted gradient echo-planar imaging (EPI) sequence (TR=2000ms, TE=28ms, FOV=240mm^2^, voxel size=3 × 3 × 3mm^3^, flip angle = 79°). Data preprocessing was performed using fMRIPrep version 1.1.4 and additional voxelwise nuisance signal removal was performed using publicly-available scripts (https://github.com/arielletambini/denoiser) to clean the data. Specifically, we denoised the data for 10 fMRIPrep-derived confounds: CSF, white matter, standardized DVARS, framewise displacement (over 0.5 mm), and six motion parameters. Functional data were high-pass filtered with a cutoff of 100 seconds, spatially smoothed with a 5 mm full-width-at-half-maximum (FWHM) Gaussian kernel, and grand-mean intensity normalized. FSL FILM pre-whitening was carried out for autocorrelation correction. First level models were run for each participant using FSL FEAT (www.fmrib.ox.ac.uk/fsl) treating the three runs as a second-level fixed effect. Events were convolved with a double-gamma hemodynamic response function. A general linear model was fit to the data with (1) a regressor for the mean signal over the duration of the cue for gain trials ($1, $2, and $3 with any probability), (2) a regressor for the mean signal over the duration of the cue for neutral trials ($0 with any probability), (3) a regressor for the mean signal over the duration of the outcome for hits, and (4) a regressor for the mean signal over the duration of the outcome for misses. We included additional temporal derivative regressors for each regressor of interest in the GLM.

#### Rutgers Shared Social Rewards

In this dataset, pairs of participants played a card-guessing game for monetary rewards in which they shared the earned outcomes of the game with three different partners––a close friend, a stranger (confederate) they met at the scan session, and a computer (*56*). All participants provided informed consent and the study was approved by the IRB at Rutgers University and the University of Medicine and Dentistry of New Jersey. One participant was scanned (N = 20, mean age=20.5, sd=2.2, females=50%) while their partners (friend, stranger) took part in the task behaviorally from the scanner control room. Participant pairs took turns playing or observing each other guess whether the value of a playing card was higher or lower than the number 5. Correct guesses resulted in a shared monetary gain of $4 for the pair on a given trial (i.e., MRI participant + friend; MRI participant + stranger); incorrect guesses resulted in a shared monetary loss of $2. Imaging data were acquired on a Siemens 3T Allegra head only scanner using a single shot gradient echo EPI sequence (TR=2000ms, TE=25ms, FOV=192, flip angle=80°, voxel size=3 × 3 × 3 mm^3^). Functional data were preprocessed using fMRIprep, including motion correction, skull-stripping, coregistration to anatomical data, realignment and normalization. Data were spatially smoothed with a 6mm FWHM Gaussian kernel and first level analyses were performed using FEAT in FSL (v 6.0.4). A GLM was computed including regressors modeling the choice (3 regressors) and outcome phases (6 regressors) of the card game for each partner. Regressors of no interest were included modeling the choice and outcome phases of missed trials. Confound regressors modeling trial-by-trial framewise displacement, head motion using realignment parameters, the first six principal components derived from aCompCor capturing physiological noise and cosine basis functions were included for each participant in each run. Whole brain beta maps for each regressor of interest in the outcome phase (i.e., monetary gain/loss with each partner) were computed.

#### Rutgers Trust Game

In this dataset, participants (N = 24, mean age=21.36, sd=3.67, 14 female) played an iterated economic trust game with either a close friend, a stranger or a computer on a given trial of the task while undergoing fMRI (*57*). All participants provided informed consent and the study was approved by the IRB at Rutgers University. MRI participants played the role of the investor in a trust game and were endowed with $1.00 on each trial of the task. They could choose to either invest (i.e., trust) their money with their partner (i.e., trustee) on a given trial or keep it (i.e., distrust). Trust decisions resulted in the $1.00 investment being multiplied by a factor of three, such that the trustee received $3.00. The trustee could then decide whether to share half of this amount ($1.50) back with the investor (i.e., reciprocate) or keep it all for themselves (i.e., defect). Unbeknownst to the participants, the trustee’s responses were randomized to reciprocate or defect with a 50% probability. Here, we focused on the outcome phase of the task on trials in which the MRI participants chose to invest with the trustees so we could examine responses to social reward (i.e., reciprocate outcomes) and social loss (i.e., defect outcomes). Imaging data were collected on a Siemens 3T Magnetom Trio whole-body scanner using a single shot gradient EPI sequence (TR=2000ms, TE=30ms, FOV=192mm, flip angle=90°, voxel size=3 × 3 × 3mm^3^). Data were preprocessed using custom scripts (https://github.com/rordenlab/spmScripts) for SPM12 and FSL (v5.09; FMRIB). Standard preprocessing was performed in SPM (motion correction, brain extraction and coregistration, slice time correction). Motion artifact was removed using ICA-AROMA in FSL(*92*). Functional data were smoothed using a 5mm kernel in FSL. We modeled each experimental condition as a separate boxcar regressor convolved with a double gamma HRF using a general linear model (GLM). The GLM resulted in whole-brain beta maps for each regressor of interest (i.e., reciprocate, defect) in the outcome phase for each partner (friend, stranger, computer). We then averaged the beta maps across partner types.

#### Neurosynth Gene Expression

This dataset is from a study that attempted to map genes to cognitive processes based on shared spatial patterns distributed throughout the brain (*50*, *51*). Gene data comes from the Allen Human Brain Atlas (AHBA)(*52*), which is a brain-wide gene expression atlas derived from transcriptome-wide microarray assessments of human brain tissue from 3,702 samples from 6 postmortem donor brains (http://www.brain-map.org). Several preprocessing steps were required to produce gene expression maps suitable for comparison with functional neuroimaging data. First, to increase measurement reliability and reduce the number of comparisons, the original authors averaged the normalized values over all probesets associated with each gene (for genes associated with more than one probeset). Second, for each individual gene and each individual donor, gene expression values across all available samples were standardized (i.e., z-scored) to remove donor wide variations in mean gene expression levels. Third, the coordinates of all microarray samples provided in the AHBA dataset were transformed into MNI152 stereotactic space. The data were spatially smooth by multiplying each point by a hard sphere of 6 mm radius centered on the microarray locations. Lastly, for each gene, a single map of brain-wide gene expression was created by averaging across all 6 of the individual donor maps. To increase reliability, we only included voxels in which gene expression levels reflected a minimum of 4 samples. All preprocessed gene maps are available on the Neurosynth website (https://neurosynth.org/). We were specifically interested in comparing the spatial patterns from the Reward Model with dopamine receptors D1, D2, D3, D4, & D5. We computed the spatial similarity using Pearson correlations.

#### Vanderbilt PET [18F]fallypride

This dataset was taken from a previously published study assessing individual differences in dopamine D2-like receptor availability and neural representations of subjective value (*53*). As part of this study, participants (N=25, mean age=20.9, sd=1.83, 52% female) underwent a PET scan with the high affinity D2/3 receptor tracer [18F]fallypride. [18 F]fallypride, (S)-N-[(1-allyl-2-pyrrolidinyl)methyl]−5-(3[18 F]fluoropropyl)−2,3-dimethoxybenzamide, was produced in the radiochemistry laboratory attached to the PET unit at Vanderbilt University Medical Center, following synthesis and quality control procedures described in US Food and Drug Administration IND 47,245. PET data were collected on a GE Discovery STE (DSTE) PET scanner (General Electric Healthcare, Chicago, IL, USA). The scanner had an axial resolution of 4 mm and in-plane resolution of 4.5 to 5.5 mm FWHM at the center of the field of view. Serial scan acquisition was started simultaneously with a 5.0 mCi (185 MBq) slow bolus injection of [18F]fallypride. CT scans were collected for attenuation correction prior to each of the three emission scans, which together lasted approximately 3.5 hours with two breaks for participant comfort. The 3 emission scans were merged temporally and motion corrected. Voxelwise images of receptor availability or binding potential (BP_ND_) were quantified using the simplified reference tissue model (*93*, *94*) in PMOD Biomedical Imaging Quantification software (PMOD Technologies, Switzerland) with the putamen as a receptor-rich region and the cerebellum as the reference region. Binding potential images represent the ratio of the specifically bound ligand ([18F]fallypride in this study) to its free concentration. All participants provided informed consent and study procedures were approved by the Institutional Review Board at Vanderbilt University. Pattern similarity of the Reward Model was computed by taking the Pearson correlation with each participant's binding potential images. We performed a signed permutation test with 5,000 iterations using the nltools toolbox (*90*).

#### Sweet Taste Dataset

This dataset examined neural responses to carbohydrate rewards (*59*). Participants (N=15, mean age=24.33 (3.92), female=47%) consumed 10 differently flavored non-caloric beverages containing 0.1% (w/v) citric and 0.0078% sucralose dissolved in demineralized water. Participants selected their 5 favorite flavors and each flavor was paired with a specific nutrient dose by adding different amounts of maltodextrin (0, 37.5, 75, 112.5, or 150 calories), with the highest dose equivalent to 12 fl oz can of soda. The five beverages along with a flavorless control solution was delivered as 1ml over 4s from syringe points with a gustometer system. Each beverage was delivered 18 times over the course of the imaging session. All participants provided informed consent on protocols approved by the Yale University School of Medicine Human Investigation Committee. Imaging data were acquired on a Siemens 3T Tim Trio scanner using a susceptibility-weighted single-shot echo planar sequence (TR=2000ms, TE=20ms, flip angle=90°, FOV=220mm, matrix 64 × 64mm, slice thickness 3 mm, 40 interleaved slices). This dataset was accessed from OpenNeuro (*85*) from (https://openneuro.org/datasets/ds000229/versions/00001) and preprocessed using fMRIPrep (*89*). Data were smoothed using a 6mm FWHM Gaussian kernel. First level single-trial models were run using nltools (*90*) using a design matrix that included boxcar regressors for each experimental condition convolved with a double gamma HRF, a linear trend, the effects of motion estimated during the realignment step using an expanded set of 24 motion parameters (six demeaned realignment parameters, their squares, their derivatives, and their squared derivatives), motion spikes between successive TRs, and global signal-intensity spikes greater than three SDs above the mean intensity between successive TRs, and a high pass filter of 120s.

#### Milkshake Dataset

Participants (N=160, mean age = 15.3, sd=1.1, 51% female) were scanned while completing a food reward fMRI paradigm(*58*), in which they viewed cues of two images (glasses of milkshake and water) that signaled impending delivery of either 0.5ml of milkshake or tasteless solution delivered using programmable syringe pumps. Participants viewed 50 cues for each stimuli and consumed 30 trials of each beverage. All participants and parents provided written informed consent to participate in this project in accordance with the IRB. Imaging data were acquired on a Siemens Allegra 3T head-only MRI scanner using T2*-weighted gradient single-shot echo planar imaging sequence (TE=30ms, TR=2000, flip angle=80°, 64 × 64 matrix, 192 ×192mm^2^ FOV, 3 × 3 × 4mm^3^ voxels). Imaging data were preprocessed and analyzed using SPM12, using DARTEL normalization, slice timing correction, unwarping using field maps, and smoothed with a 6mm FWHM Gaussian kernel. First level models included box car regressors for each experimental condition, realignment parameters, spikes identified using ART. First level contrast images aggregating runs for each condition were shared by the first and senior authors.

#### UCLA Balloon Analog Risk Task

We used the Balloon Analog Risk Task (BART) to probe psychological processes related to making risky decisions. In this task, participants (N=124, mean age=31.58, sd= 8.81, female = 48%) were allowed to pump a series of virtual green (risky) and white (safety) balloons. On each trial, participants chose to either pump the balloon or cash out and collect their accumulated earnings for that round. For the risky balloons, a successful pump yielded 5 points and the participant was given the opportunity to continue to pump or cash out (a maximum of 12 pumps were possible). An unsuccessful pump led to the balloon exploding and the participant earned no points for the round. Balloons exploded randomly based on a random draw from a uniform distribution over numbers of pumps. In the safe condition, participants could pump, but the balloon never exploded, nor did the participant win any points. This data was collected by the Consortium for Neuropsychiatric Phenomics as part of a larger study focused on understanding the dimensional structure of cognitive processing in healthy individuals and those diagnosed with neuropsychiatric disorders (*65*). All participants gave written informed consent following procedures approved by the Institutional Review Boards at UCLA and the Los Angeles County Department of Mental Health. Imaging data were collected on a Siemens Trio 3T scanner using EPI sequence (TR=2000ms, TE=30ms, flip angle=90°, matrix=64 × 64, FOV=192mm, 34 oblique slices). Data was accessed from OpenNeuro (https://openneuro.org/datasets/ds000030/versions/1.0.0) and preprocessed using fMRIPrep (*89*). First level single-trial models were run using nltools (*90*). Data were smoothed using a 6mm FWHM Gaussian kernel. Each trial was convolved with a double gamma HRF and and additional covariates included a linear trend, the effects of motion estimated during the realignment step using an expanded set of 24 motion parameters (six demeaned realignment parameters, their squares, their derivatives, and their squared derivatives), motion spikes between successive TRs, and global signal-intensity spikes greater than three SDs above the mean intensity between successive TRs, and a high pass filter of 120s.

#### HCP Social Task

In the HCP social task, participants (N=484, mean age = 29.24, 59% female) watched short video clips of objects (e.g., squares, circles, triangles) either interacting with each other or moving randomly (*67*, *95*). Participants viewed five 20s clips of each condition, which were separated by 15s fixation blocks. After each video clip, participants reported whether they believed the objects were interacting or not. The data acquisition, preprocessing, and first level models for this task were identical to protocol described in the reward task. The beta estimates for the average response to each condition were accessed via the OpenAccess AWS S3 bucket provided by the HCP.

#### Self Referential

In the self-referential task, participants were Chinese graduate students who recently arrived in the United States (N=27, mean age=24.11, 48% female). Participants completed a trait-judgment task (*68*, *69*), in which they made three different types of judgments: a self-judgment (i.e. does optimistic describe you?), a mother-judgment (i.e., does optimistic describe your mother?), and a font-judgment (i.e. is this word printed in bold-faced letters?) condition presented in either Mandarin or English. The current study only used the trials presented in Mandarin. All participants provided informed consent in accordance with the guidelines set by the Committee for the Protection of Human Subjects at Dartmouth College. Imaging data were collected using a Philips Intera Achieva 3T scanner and a 32 channel head coil using an EPI sequence (TR=2,500ms, TE=35ms, flip angle=90, FOV=240mm, voxel size=3 × 3 × 3 mm^3^). Data were preprocessed using SPM8, which included slice timing correction, unwarping, realignment, motion correction, normalization, and spatial smoothing with a 6 mm FWHM Gaussian kernel. For each participant, the three judgment conditions were modeled separately in the GLM with a boxcar regressor convolved with a double gamma HRF. Additional covariates of no interest included a linear trend, and 6 realignment parameters. We used beta images containing average activity when participants made self-judgments and compared this to activity elicited by trials in which participants made judgments about the font.

#### Boulder Pain

In this study (*37*, *38*), participants (N=28, mean age=25.2, sd= 7.4, female = 40%) received thermal pain stimulation applied to the volar surface of the left forearm and dorsal surface of the left foot using a TSA-II Neurosensory Analyzer (Medoc Ltd., Chapel Hill, NC) with a 16 mm Peltier thermode end plate. All participants consented to the experimental procedure which was approved by the University of Colorado IRB. Three levels of thermal stimulation were applied to four different locations on both the upper limb (i.e., volar surface of the left forearm) and lower limb (i.e., dorsal surface of the left foot) at low (46°C), medium (47°C), and high (48°C) intensities for 11 seconds while participants were being scanned on a Siemens Tim Trio 3T MRI scanner. Functional images were acquired with an EPI sequence (TR = 1300 ms, TE = 25 ms, field of view = 220 mm, 64 × 64 matrix, 3.4 × 3.4 × 3.4 mm voxels, 26 interleaved slices with ascending acquisition, parallel imaging with an iPAT acceleration of 2). Data were accessed from NeuroVault (*35*) (https://neurovault.org/collections/504/). All images were preprocessed by the original authors using SPM8 (Wellcome Trust Centre for Neuroimaging, London, UK) and custom Matlab functions. Functional images were corrected for slice-acquisition-timing and motion using SPM8. They were then warped to SPM’s normative atlas using warping parameters estimated from coregistered, high-resolution structural images, interpolated to 2 × 2 × 2 mm voxels, and smoothed with an 8 mm FWHM Gaussian kernel. Prior to preprocessing of functional images, global outlier time points (i.e., “spikes”) were identified by computing both the mean and the standard deviation (across voxels) of values for each image for all slices. Mahalanobis distances for the matrix of slicewise mean and standard deviation values (concatenated) were computed for all functional volumes (time), and any values with a significant χ^2^ value (corrected for multiple comparisons) were considered outliers (less than 1% of images were outliers). The output of this procedure was later used as a covariate in the first level models. First-level GLM analyses were conducted in SPM8. Boxcar regressors, convolved with the canonical hemodynamic response function, were constructed to model periods for the 2 sec cue presentation, the 5, 7, or 11 sec variable prestimulus fixation period, the 11 sec thermal stimulation, and the 4 sec rating periods. Additional covariates included a high-pass filter of 224 sec, outliers, and 24 expanded motion parameters (6 realignment, their derivatives, squares, and squared derivatives). Contrast maps were created for the low, medium, and high stimulation periods collapsing across cues (i.e., low, medium, and high) and body site (i.e., upper limb and lower limb).

#### Pittsburgh IAPS

Participants (N=182, mean age = 42.77, sd = 7.3, female = 52%) were recruited from the greater Pittsburgh area to complete an affective reappraisal task (*38*). All participants gave informed consent in accordance with the guidelines set by the IRB at The University of Pittsburgh. Participants viewed 15 negative photographs and 15 neutral photographs selected from the International Affective Picture System (IAPS) (*96*) and were instructed to either (a) “look” and maintain their attention to the photos when they came on screen and allow their emotional reactions to occur naturally or (b) “decrease” and change the way they thought about the image to feel less negative (see (*38*, *97*) for full task and IAPS stimulus details). Data were collected on a Siemens 3T Trio TIM whole-body scanner (TR=2000ms, TE=29ms, Flip Angle=150, FOV=200mm, matrix=64 × 64, 34 3mm slices). The data were accessed via Neurovault (*35*) at (https://neurovault.org/collections/503/). Data was preprocessed by the original authors using SPM8, including unwarping, realignment, coregistration, normalization, spatial smoothing with a 6 mm FWHM Gaussian kernel and high pass filtering (180 sec cutoff). Five separate regressors indicating different rating levels (1 to 5) were modeled in the GLM for each participant as well as 24 covariate regressors modeled movement effects (6 realignment parameters demeaned, their 1st derivatives, and the squares of these 12 regressors). In this study, we used single trial responses from only the look condition, which were presented for 7 seconds and only included participants (N = 93) who rated images either a 1 (neutral) or a 5 (most negative).

#### HCP Language

In the HCP language task (*98*), participants (N=482, mean age = 29.24, 59% female) listened to eight blocks of approximately 30 s stories adapted from Aesop’s stories (5-9 sentences each). These blocks were interleaved with eight 30s blocks of math problems which included a series of arithmetic operations that were also presented auditorily. After each block, participants were instructed to complete a forced choice test demonstrating semantic understanding of the content. The data acquisition, preprocessing, and first level models for this task were identical to protocol described in the reward task. The beta estimates for the average response to each condition were accessed via the OpenAccess AWS S3 bucket provided by the HCP.

#### HCP Working Memory

In the HCP working memory task, participants (N=493, mean age = 29.24, 59% female) completed a version of the N-back task (*99*, *100*), in which participants were presented visual stimuli from four separate categories (i.e., faces, places, tools, and body parts) that have previously been shown to engage distinct cortical areas. Participants were instructed to complete two separate tasks across 16 blocks of stimuli (8 blocks per task). Each block contained 10 images presented for 2.5s each and contained two “targets” and 2-3 “lures”. For the “2-Back” task, participants were instructed to press a button whenever the current “target” stimulus was the same one as two images back. The lures were also a repeated image, but were repeated one or three images back. This task is meant to probe active working memory maintenance. For the “0-Back” task, participants were presented with a “target” image at the beginning of the block and were instructed to press a button every time the stimulus is presented again at the end of the block. This task controls for goal maintenance and motor responses, but does not contain an active working memory load component. The data acquisition, preprocessing, and first level models for this task were identical to protocol described in the reward task. The beta estimates for the average response to each condition were accessed via the OpenAccess AWS S3 bucket provided by the HCP.

#### Stop Signal Task

In this study, participants (N=19, mean age=23.75, sd=5.87, 47% female) completed a manual stop signal task (SST) (Neurovault task001) (*101*). For the go trials in this task, participants were asked to press on the right or left button according to whether the letter “T” or “D” was shown on the screen. For stop trials, an auditory tone cue signaling stop was played after the letter being shown with some delay (stop-signal delay; SSD), and participants were asked to inhibit their approaching responses toward the button. Throughout the task, the length of SSD changed according to whether participants succeeded or failed to inhibit their responses in order to maintain the accuracy rate at 50%. All participants gave informed consent according to a procedure approved by the University of California Los Angeles Human Subject Committee. Data were collected on a 3T Siemens Allegra MRI scanner using an EPI sequence (TR=2000ms, TE=30ms, flip angle=90°, matrix 64 ×64, FOV=200mm). This dataset was accessed from NeuroVault (*35*) from (https://neurovault.org/collections/1807/). Preprocessing was performed using FSL version 3.3 by the authors of the original study and included coregistration, realignment, motion correction, denoising using MELODIC, normalization, spatial smoothing with a 5 mm FWHM Gaussian kernel, and high-pass filtering with a 66s cutoff. Each condition (i.e., go, inhibition-success, inhibition-failure, and nuisance events) were modeled as a separate boxcar regressor and convolved with a double gamma HRF using a GLM. Temporal derivatives and 6 motion parameters were included as covariates.

#### MIT Deprivation Task

This study investigated the neural signals of social isolation (*41*). Participants (N=30, mean age=26.75, sd=5.6, female=73%) completed three separate scanning sessions (i.e., baseline condition, food deprivation, and social isolation). Across all three sessions, participants were scanned while they completed an image viewing paradigm, which included viewing 54 images within three separate conditions. In the social condition, participants viewed groups of individuals as they meet, talk, laugh, smile, etc. In the food condition, participants viewed different kinds of highly palatable foods such as cake, pizza, chocolate, etc. In the control condition, participants viewed images of attractive flowers. On each trial, participants saw a single photograph and 3-5 word verbal description, for 5 sec. The combination of visual and verbal cues was intended to maximize deep semantic processing of the relevant attributes. In the food deprivation scanning session, participants were asked to abstain from consuming any food or drinks/coffee (except water) for 10 hours before the fMRI session. They could engage in any social or non-social activities they wanted to but were asked to abstain from exercising in order to avoid exhaustion. In the social isolation condition, participants remained in a room for 10 hours and were not able to interact socially for the duration of the isolation. Participants were provided with puzzles, Sudoku, coloring pages, non-social games (e.g., Tetris, Bubble Shooter, etc.) and drawing/writing supplies. Participants were able to eat any food they wanted during isolation. In the baseline condition, participants did not undergo any experimental manipulation before scanning. All participants consented to participate in the experiment in accordance with MIT’s institutional review board.

The stimuli for the image viewing task were tailored to each individual's preferred foods and modes of social interaction. During the initial screening, participants were asked to list their top ten favorite foods and social activities. Stock photographs illustrating these specific foods and activities were selected from a large public database (https://www.pexels.com/), and then verbal labels were added using the participant’s own descriptions. Food descriptions included “fluffy syrup-drenched pancakes”, “creamy cheesy macaroni”, “refreshing mixed fruit salad”, and “yummy vanilla cake with sprinkles”. Social descriptions included “chatting and laughing together”, “joking around with friends”, “supporting each other through workouts”, “enjoying a conversation together.” Social pictures were all matched for gender of participants (i.e., for a male participant, all social photographs included at least one man). Control trials presented attractive photographs of flowers accompanied by positive valence verbal descriptions.

Data were acquired on a Siemens Prisma (TR = 2,000 ms, TE = 30 ms, FOV = 210 mm, 70 × 70 matrix, yielding a voxel size of 3 × 3 × 3 mm^3^) acquired as a partial-head volume in an anteroposterior phase-encoding direction using interleaved slices and were accessed from OpenNeuro (*85*) (https://openneuro.org/datasets/ds003242/versions/1.0.0). Data were preprocessed using fMRIPREP (*89*) and smoothed using a 6mm FWHM Gaussian filter. First level single-trial models were run using nltools (*90*). Each trial was convolved with a double gamma HRF and additional covariates included a linear trend, the effects of motion estimated during the realignment step using an expanded set of 24 motion parameters (six demeaned realignment parameters, their squares, their derivatives, and their squared derivatives), motion spikes between successive TRs, and global signal-intensity spikes greater than three SDs above the mean intensity between successive TRs, and a high pass filter of 120s.

#### Mixed Gamble Task

This study investigated the neural correlates of loss aversion while individuals decided whether to accept or reject gambles that offered a 50% chance of gaining or losing money. All participants were free of neurological and psychiatric history and gave informed consent to participate according to a protocol approved by the University of California, Los Angeles Institutional Review Board. Participants (N=16, mean age=22, sd=2.9, 56% female) made decisions across 85 trials that varied potential gain amounts between $10 and $40 and potential losses between $5 and $20 in order to create a variety of situations at and around the point where the potential gain is twice the loss. Gains and losses varied across trials, but gains were designed to be twice as high as losses. This was designed to create a range of subject reactions from strong rejection to strong acceptance. Participants were paid at the end of the experiment for one randomly selected trial to ensure incentive compatibility. Data were acquired using a 3T Siemens AG Allegra MRI scanner using an EPI sequence (TR=2000ms, TE=30ms, flip angle=90, FOV=200mm, matrix=64 × 64). Data were accessed from OpenNeuro (https://openneuro.org/datasets/ds000005/versions/00001) and preprocessed using fMRIPrep (*89*). Data were smoothed using a 6mm FWHM Gaussian kernel. First level single-trial models were run using nltools (*90*). Each trial was convolved with a double gamma HRF and and additional covariates included a linear trend, the effects of motion estimated during the realignment step using an expanded set of 24 motion parameters (six demeaned realignment parameters, their squares, their derivatives, and their squared derivatives), motion spikes between successive TRs, and global signal-intensity spikes greater than three SDs above the mean intensity between successive TRs, and a high pass filter of 120s. We computed the pattern similarity between the Reward Model and each trial for each participant using Pearson correlations.

#### Naturalistic Viewing Task (FNL)

In this study, participants (N=35, mean age = 19.0, sd=1.07, 74% female) were recruited to watch the pilot episode of the character driven television drama, Friday Night Lights (FNL) while being continuously scanned with fMRI (*4*). All participants provided informed consent in accordance with a protocol approved by the Committee for the Protection of Human Subjects at Dartmouth College. Data were acquired using a 3T Siemens Magnetom Prisma scanner (Siemens, Erlangen, Germany) with a 32-channel head coil (TR/TE = 2000/25 ms, flip angle = 75°, resolution = 3 mm3 isotropic voxels, matrix size = 80 by 80, and FOV = 240 mm by 240 mm, GRAPPA=2) for 45 minutes (1364 TRs). All data are available on OpenNeuro (https://openneuro.org/datasets/ds003521/versions/1.0.0). Data underwent a standard preprocessing pipeline https://github.com/cosanlab/cosanlab_preproc, which included motion correction, nonlinear spatial normalization, spatial smoothing using a 6mm FWHM Gaussian kernel. Data were denoised using a voxel-wise GLM to remove variance associated with the mean, linear, and quadratic trends, mean activity from a cerebral spinal fluid mask, the effects of motion estimated during the realignment step using an expanded set of 24 motion parameters (six demeaned realignment parameters, their squares, their derivatives, and their squared derivatives), motion spikes between successive TRs, and global signal-intensity spikes greater than three SDs above the mean intensity between successive TRs. We applied the Reward Model to the entire preprocessed and denoised time series for each participant.

#### Facial Expressions

Participants (N=20, mean age=18.9, sd=0.91, 65% female) were recruited from the Department of Psychological Brain Sciences at Dartmouth College to watch the first four episodes of the first season of FNL over two separate 2-hour sessions. Here, we report results from episode one. All participants provided informed consent in accordance with a protocol approved by the Committee for the Protection of Human Subjects at Dartmouth College. Facial expressions during the experiment were monitored using GoPro HERO 4 cameras recording at 120 frames/s at 1920 by 1080 resolution. Each camera was positioned using a custom facecam headsets developed by our group (*76*). This approach is invariant to head motion and minimizes many types of facial occlusions. Recorded videos were then temporally aligned to the episodes by minimizing differences in audio intensity using our open source Python FaceSync toolbox version 0.0.8 (*76*). Facial behavioral features consisting of 20 facial AUs, a standard for measuring facial muscle movement based on the Facial Action Coding System (*77*), were extracted using FACET^®^ (*102*) accessed through the iMotions biometric research platform. Data were downsampled to 0.5 Hz. We used our Python Facial Expression Analysis Toolbox (Py-Feat) version 0.4 to visualize the facial morphometry of our Face Model using min-max feature scaling along the interval of [0,1] (*78*). See the original paper for full details about image acquisition, preprocessing, and denoising procedures (*4*, *103*). Data are available to be downloaded from OSF (https://osf.io/f9gyd/).

### Model Training

We trained our model using data from the Delgado Gambling task collected as part of the Human Connectome Project made available via an AWS S3 bucket (N=490). We randomly selected 80% of the participants to serve as training data (N=392) and 20% of participants to serve as a separate hold out test dataset (N=98). For the training data, we tried to remove as much individual subject variability as possible by subtracting the subject mean out of each map and standardizing the data within each image. We trained a linear Support Vector Machine (SVM) to classify reward trials from punishment trials using 5-fold cross-validation procedure where data from the same subject was held out together. This allowed us to estimate the generalizability of this model to new participants that have been preprocessed in a similar manner. We performed a more rigorous validation using the separate hold out set. Unlike the cross-validated analyses, we used the full training dataset to train the model (n=392) and only tested the model once on the hold out test dataset (n=98).

To establish the face validity of our model, we used a parametric bootstrap to identify which voxels most reliably contributed to the classification, which involved retraining the model 5,000 times after randomly sampling participants with replacement and thresholding at FDR q < 0.0001. This procedure is purely for visualization to establish the face validity of the model and not used for spatial feature selection (*44*).

This bootstrap procedure also allowed us to to estimate the consistency of the spatial pattern across different random samples of the data. We computed the pairwise spatial similarity of the whole-brain pattern estimated across each bootstrap iteration and observed a high level of spatial consistency, r=0.93, p < 0.001 (*38*). While this procedure allows us to estimate the variation in the model weights across different subsamples of participants from the full training dataset, it is important to note that this estimate is likely slightly inflated compared to a more traditional reliability estimate as the data are not fully independent across bootstraps.

### Model Testing

#### Forced Choice Accuracy

To evaluate the convergent and discriminant validity of the Reward Model to other psychological constructs, we tested our reward classification models using forced choice accuracy tests. Forced choice tests compare the relative spatial similarity of each brain map to the Reward Model within the same participant using Pearson correlations and then select the map with the overall highest similarity. We performed hypothesis tests using a permutation approach, by randomly permuting the order of the images for each participant 10,000 times to generate a null distribution, and count the number of instances in which our average accuracy across participants exceeds this null distribution. We were only interested in whether the target condition was significantly greater than the reference condition, so we report one-tailed tests. Forced choice tests are well suited for fMRI because they do not require signals to be on the same measurement scale across individuals or scanners (*30*) and have an interesting property in that they are equivalent to sensitivity, specificity, and area under the ROC curve (AUC).

#### Virtual Lesion

We performed a virtual lesion analysis (*38*) to determine how weights in regions outside the core reward network contributed to the overall prediction. This required training a new predictive model after performing spatial feature selection to remove the contribution of voxels outside regions traditionally considered to be involved in reward processing. We identified voxels that are frequently implied by the term “reward” using Neurosynth (*28*). Neurosynth uses a database containing 14,371 neuroimaging studies and computes a bag of words to represent the frequency of words used in each paper. It uses a naive bayes classifier with a uniform prior to determine the likelihood of the term given the pattern of activations reported in the paper. We created a binary mask using the neurosynth reverse inference maps thresholded at FDR 0.01. We further smoothed the mask by applying a 5mm Gaussian kernel and thresholded the map to include voxels that exceeded a z-score of 3.5. These values were arbitrarily selected and produce a mask that includes contiguous core regions of the reward network (e.g., ventral striatum, SN/VTA, vmPFC). We used this mask to select 7,322 voxels from the 238,955 included in the whole brain model and retrained the Reward Model using the identical procedure described above. Importantly, this procedure removes the influence of regions that may be peripheral to reward processing (e.g., sensory cortex). We next performed nested model comparisons of the predictions of the full whole-brain model compared to the sparse masked model in order to determine the impact of performing a virtual lesion or ablating weights outside core reward regions. This entailed using a mixed logistic regression using the pymer python wrapper (*104*) around the R lme4 package (*105*) to predict the classification accuracy of each subject based on the type of model indicated by a dummy code. This analysis was performed separately for each dataset. Participant IDs were used as a random intercept and slope. The results of this virtual lesion analysis for each dataset using forced choice accuracy are reported in Table S1. Positive z-values indicate that the whole-brain model yielded a higher accuracy relative to the nested sparse model, while negative values indicate that the accuracy increased in the sparse model.

We openly share both the whole-brain and the sparse Reward Models and recommend that these models be used interchangeably depending on the researcher’s goals. The whole-brain model appears to be more sensitive to detecting rewards when the task is similar to the original training task. The sparse model appears to be more specific, but may miss important information represented in other cortical regions.

#### Loss Aversion

We used a linear model of prospect theory to estimate each participant’s individual loss-aversion parameter. The original Prospect Theory model proposed a nonlinear weighting function (*79*), here we simplify the model by assuming equal decision weights for a 0.5 probability to gain or lose money as proposed in (*42*), 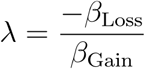. We use a mixed effects logistic regression using lme4 (*105*) using the pymer python wrapper (*104*) to predict decisions to accept or reject the gamble based on the possible gains or losses for that trial treating participant IDs as a random intercept and slope. We used the subject level BLUPs from this model *β*_Loss_ and *β*_Gain_ to estimate λ. We used a similar approach to estimate each participant’s neural loss-aversion parameter λ. We computed the pattern similarity of our Reward Model with single trial beta estimates of each trial, and then predicted variation in the Reward Model predictions using the magnitude of loss and gains associated with each trial. Similar to the behavioral estimation λ, we estimated each individual participant’s loss aversion parameter using the subject level BLUPs from the regression model *β*_Loss_ and *β*_Gain_. We used a Spearman *ρ* rank correlation to estimate the magnitude of the relationship between the behavior and brain loss-aversion parameters and a permutation test with 10,000 to perform a hypothesis test. We note that *β*_Loss_ and *β*_Gain_ estimated from predicting the Reward Model pattern similarity were very small, so the values we report in the results section are standardized estimates.

#### Intersubject Correlation (ISC)

For the naturalistic image viewing task, we used intersubject correlation (ISC) to examine the reliability of neural dynamics in response to a dynamic stimulus across individuals (*74*, *75*, *106*). We computed the pairwise correlation between participants’ predicted time series after taking the spatial similarity with each image and the Reward Model. We computed the mean of the lower triangle of the subject by subject-by-subject correlation matrix after performing a Fisher R to Z transformation and then inverted the transformation after computing the mean. To perform inferences if there was an overall significant level of synchronization, we used the subject-wise bootstrap procedure proposed by (*107*) with 5,000 bootstraps using the nltools software package (*90*). See (*75*, *108*) for a tutorial on ISC.

#### Face Expression Model

To create the facial expression model, we used the standardized average predicted AU response that was downsampled to 0.5Hz to match the neuroimaging data. We used a mixed effects regression model using the pymer4 toolbox (*104*) to predict the subject-specific pattern similarity time series after applying the Reward Model to the naturalistic FNL fMRI data. We used min-max feature scaling to scale the *β*_*AU*_ estimates from the regression model to lie on the interval of [0,1]. We visualized the weights of the model using the py-feat toolbox (*78*). To assess the predictive accuracy of the model, we used 5-fold cross-validation where data from the same subject was held out together to train the model on a subset of the data and then assessed the ability of the model to account for changes in each individual fMRI participants’ predicted reward responses using a Spearman *ρ* rank correlation. We used a sign permutation test with 5,000 permutations to perform inference over participants. This procedure provides an unbiased estimate of how well the model could account for the fluctuating dynamics of the Reward Model predictions over the sequence of events in the final football scene in the television episode.

## Acknowledgments

This research was supported by funding from the National Institute of Health (R01MH116026 and R56MH080716 to L.J.C.; R15MH122927 to D.S.F.; R01MH084081 to M.R.D., R01AG043458 to G.R.S.-L. & D.H.Z, and R21DA033611 to D.H.Z.), the National Science Foundation (NSF CAREER 1848370 to L.J.C.), the Young Scholar Fellowship Program by the Ministry of Science and Technology and funding from National Science and Technology Council in Taiwan (MOST 109-2636-H-002-006, 110-2636-H-002-004, 111-2628-H-002-004, and NSTC 111-2423-H-002-008-MY4 to P.-H.C.), as well as the Cambridge Philosophical Society (Henslow Research Fellowship to L.T.). All of the code used to perform the analysis in the paper is available on github https://github.com/cosanlab/reward_signature. The Reward Models will be uploaded to Neurovault pending publication of this manuscript. We thank Drs. Sonja Yokum and Eric Stice for generously sharing their milkshake dataset with us, all of the authors that publicly shared their datasets, and the OpenNeuro support team for helping resolve issues we encountered accessing several of these datasets.

