## Supplemental Materials for "A neural signature of reward"

#### Figures

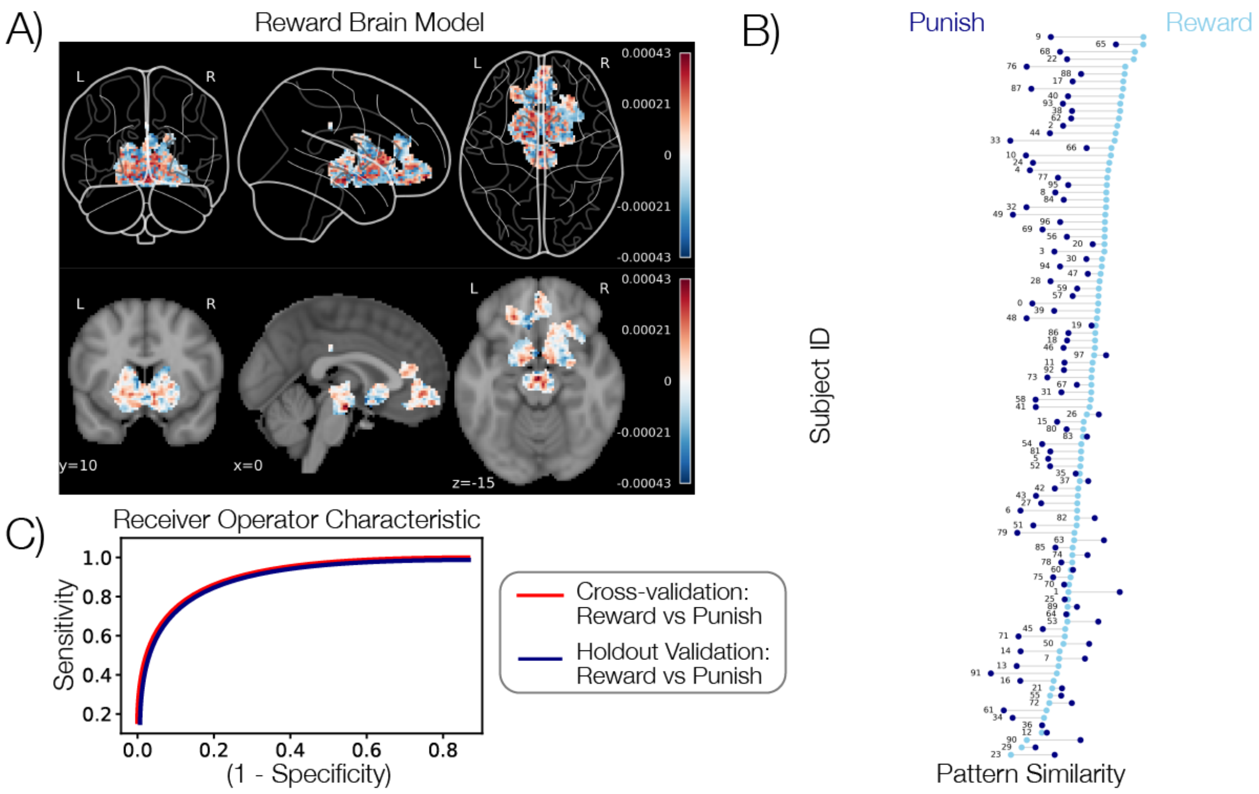


***Figure S1. Sparse Reward Model training and validation****. Panel A depicts the pattern weights for the masked sparse Reward Model using the full training dataset (N=392). Positive weights indicate an increased likelihood of classifying an image as a reward, while negative weights indicate an increased likelihood of classifying an image as a punishment. Panel B depicts the results of the forced choice accuracy tests on the separate holdout dataset (N=98). The x-axis indicates the pattern similarity to the Reward Model and each line represents a single test subject. Light blue dots indicate the reward condition while dark blue dots indicate the punishment condition. Panel C depicts the receiver operator characteristic curves for testing the Reward Model in cross-validation within the training dataset (red line, 81% forced choice accuracy) and on the separate holdout dataset (blue line, 81% forced choice accuracy).*

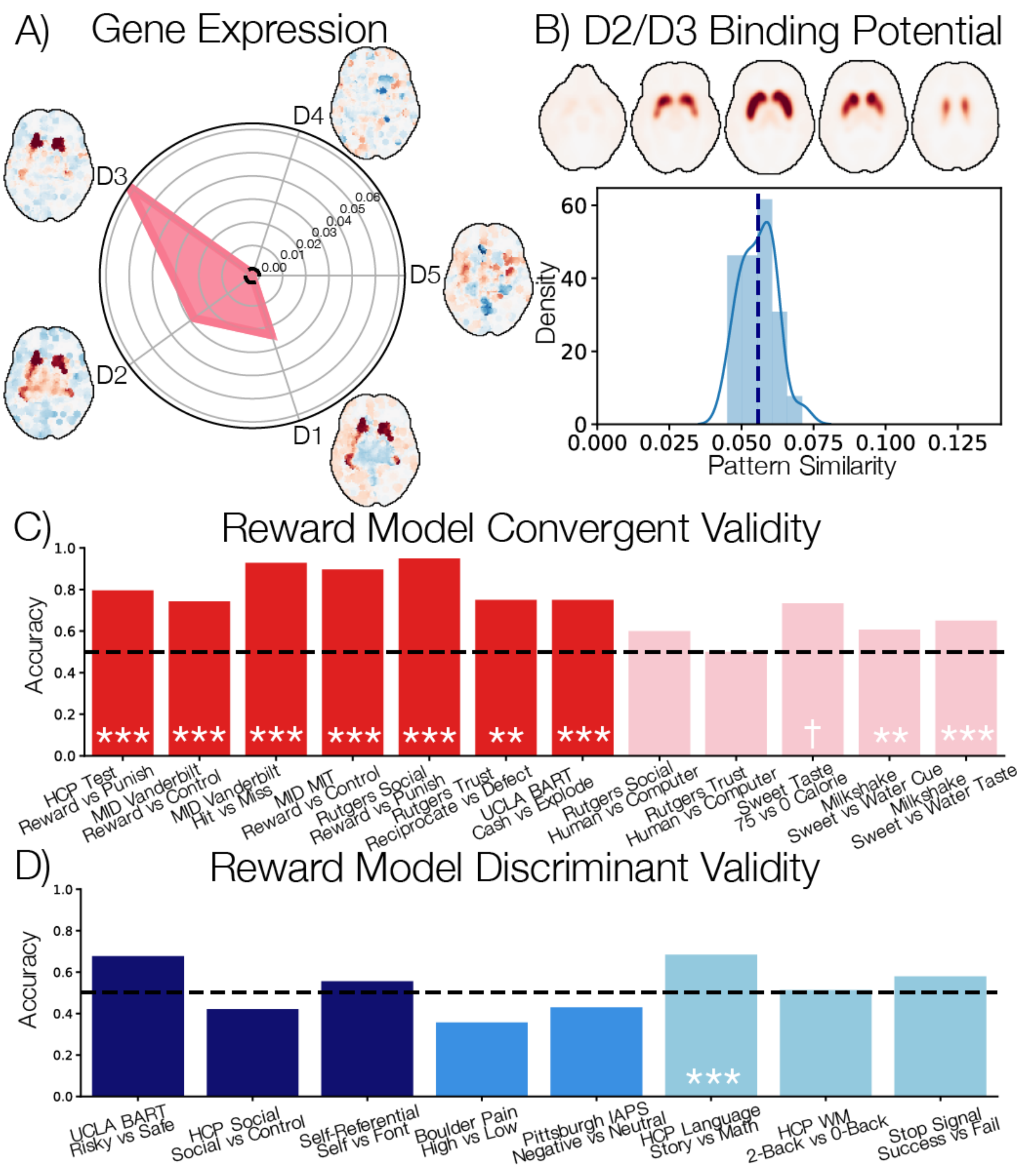


***Figure S2. Construct Validity.*** *A) Spatial similarity of the Reward Model with dopamine receptor gene expression derived from transcriptome-wide microarray assessments of brain tissue collected from 3,702 samples across 6 human donors provided by the Allen Human Brain Atlas. Values indicate Pearson correlation coefficients. B) Spatial similarity of the D2 receptor availability from participants (N=25) that underwent positron emission tomography scans (PET) using the high affinity D2-like receptor tracer*

*[18F]fallypride. C) Convergent validity of the sparse Reward Model in discriminating between the reward and no reward contrasts from multiple studies. Red values are all contrasts between conditions in which participants receive a monetary outcome compared to nothing or losing money. Pink values indicate contrasts in which participants receive other types of rewards such as social or tasting sweet drinks. Level of statistical significance is indicated by † for $p < 0.1$, ** for $p < 0.01$, & *** for $p < 0.001$. D) Divergent validity of the Reward Model in discriminating between contrasts from multiple studies. Navy values depict accuracy from risky vs safe contrasts from studies using the BART task and social and non-social contrasts. Medium blue values indicate accuracy values from negative affect studies. Light blue values indicate contrasts probing more traditional cognitive faculties such as language, working memory, and cognitive control.*

### Tables

| Study | Contrast | Sample Size | Pattern Similarity (SD) | Accuracy | Accuracy p-value | Lesion Z | Lesion p-value |
|---|---|---|---|---|---|---|---|
| HCP Reward Test | Reward vs Punish | 98 | 0.01 (0.01) | 0.81 | 0.0001 | 1.11 | 0.267 |
| MIT MID | Reward vs Neutral | 29 | 0.01 (0.01) | 0.90 | 0.0001 | 0.00 | 1.0 |
| Vanderbilt MID | Reward vs Neutral Anticipation | 70 | 0.00 (0.01) | 0.74 | 0.0002 | 4.47 | < 0.001* |
| Vanderbilt MID | Hit vs Miss Outcome | 70 | 0.01 (0.01) | 0.93 | 0.0002 | 2.11 | 0.035 |
| Rutgers Social Reward | Reward vs Punish | 20 | 0.01 (0.01) | 0.95 | 0.0001 | 0.41 | 0.68 |
| Rutgers Social Reward | Friend + Stranger Reward vs Computer Reward | 20 | 0.00 (0.00) | 0.60 | 0.2537 | 0.34 | 0.736 |
| Rutgers Trust | Reciprocate vs Defect | 24 | 0.00 (0.01) | 0.75 | 0.0100 | 2.70 | 0.007* |
| Rutgers Trust | Friend + Stranger Reciprocate vs Computer Reciprocate | 24 | 0.00 (0.01) | 0.50 | 0.5919 | -0.37 | 0.712 |
| Neurosynth Gene | D1 - Gene Expression Similarity | | | 0.02 | | | |
| Neurosynth Gene | D2 - Gene Expression Similarity | | | 0.03 | | | |
| Neurosynth Gene | D3 - Gene Expression Similarity | | | 0.06 | | | |

| | | | | | | | |
|---|---|---|---|---|---|---|---|
| Neurosynth Gene | D4 - Gene Expression Similarity | | | 0.00 | | | |
| Neurosynth Gene | D5 - Gene Expression Similarity | | | 0.00 | | | |
| Vanderbilt Fallypride | D2-like Binding Similarity | 25 | | 0.06 | 0.0002 | | |
| Sweet Taste | Calorie 75 vs Calorie 0 | 15 | 0.00 (0.01) | 0.73 | 0.0575 | 0.53 | 0.598 |
| Milkshake | Milkshake Cue vs Water Cue | 160 | 0.00 (0.01) | 0.61 | 0.0055 | 1.15 | 0.248 |
| Milkshake | Milkshake Receipt vs Water Receipt | 160 | 0.00 (0.01) | 0.65 | 0.0001 | -3.06 | 0.002* |
| UCLA BART | Cash vs Explode | 124 | 0.01 (0.01) | 0.75 | 0.0001 | 6.67 | < 0.001* |
| UCLA BART | Risky vs Safe | 124 | 0.00 (0.01) | 0.68 | 0.0002 | -2.86 | 0.004* |
| HCP Social | Social vs Control | 484 | 0.00 (0.01) | 0.42 | 1.0000 | 2.46 | 0.014* |
| Self Referential | Self vs Font | 27 | 0.00 (0.00) | 0.56 | 0.3373 | -3.01 | 0.003* |
| Boulder Pain | High vs Low | 28 | 0.00 (0.01) | 0.36 | 0.9544 | 1.32 | 0.187 |
| Pittsburgh IAPS | Negative vs Neutral | 93 | 0.00 (0.01) | 0.43 | 0.9235 | 2.18 | 0.029* |
| HCP Language | Story vs Math | 482 | 0.00 (0.00) | 0.68 | 0.0001 | -11.27 | < 0.001* |
| HCP Working Memory | 2 Back vs 0 Back | 493 | 0.00 (0.01) | 0.51 | 0.2999 | -6.48 | < 0.001* |
| Stop Signal | Success vs Fail | 19 | 0.00 (0.01) | 0.58 | 0.3253 | -0.68 | 0.498 |

| | | | | | | | |
|---|---|---|---|---|---|---|---|
| MIT Deprivation | Social Deprivation - Social vs Baseline | 30 | 0.00 (0.00) | 0.70 | 0.0221 | 2.52 | 0.012* |
| MIT Deprivation | Social Deprivation - Social vs Food | 30 | 0.00 (0.00) | 0.63 | 0.0992 | 1.05 | 0.292 |
| MIT Deprivation | Food Deprivation -Food vs Baseline | 30 | 0.00 (0.00) | 0.80 | 0.0014 | 1.52 | 0.128 |
| MIT Deprivation | Food Deprivation - Food vs Social | 30 | 0.00 (0.00) | 0.67 | 0.0495 | 3.57 | < 0.001* |
| Mixed Gamble | Accept vs Reject | 16 | | 0.69 | 0.0984 | 0.40 | 0.687 |
| Friday Night Lights | Temporal ISC | 35 | | 0.05[T] | 0.0002 | | < 0.001* |

**Table S1. Convergent and Discriminant Validity of Masked Reward Signature.** We performed a virtual lesion analysis to compare the accuracy of the whole-brain Reward Model compared to the sparse masked Reward Model using a mixed effects logistic regression. We report the z-statistic for the hypothesis test on the parameter estimate and the corresponding p-value. Positive z values indicate that the whole-brain model had a higher accuracy than the sparse model. Negative values indicate the reverse. *indicates significance at $p < 0.05$. [T]indicates temporal intersubject correlation (ISC).
